# Correlative In Situ Cryo-ET Reveals Cellular and Viral Remodeling Associated with Selective HIV-1 Core Nuclear Import

**DOI:** 10.1101/2025.03.04.641496

**Authors:** Zhen Hou, Yao Shen, Stanley Fronik, Juan Shen, Jiong Shi, Jialu Xu, Long Chen, Nathan Hardenbrook, Christopher Thompson, Sarah Neumann, Alan N. Engelman, Christopher Aiken, Peijun Zhang

**Author notes:** Equal contribution (Zhen Hou, Yao Shen).

## Abstract

Lentiviruses like HIV-1 infect non-dividing cells by traversing the nuclear pore, but studying this process has been challenging due to its scarcity and dynamic nature in infected cells. Here, we developed a robust cell-permeabilization system that recapitulates HIV-1 nuclear import and established an integrated cryo-correlative workflow combining cryo-CLEM, cryo-FIB, and cryo-ET for targeted imaging of this process. These advancements enabled the successful capture of 1,899 HIV-1 cores at various stages of nuclear import. Statistical and structural analyses of native wild-type and mutant cores revealed that HIV-1 nuclear import depends on both capsid elasticity and nuclear pore adaptability, as well as nuclear factors such as CPSF6. Brittle cores fail to enter the nuclear pore complex (NPC), while CPSF6-binding-deficient cores stall inside the NPC, resulting in impaired nuclear import. Intriguingly, nuclear pores function as selective filters favoring the import of smaller, tube-shaped cores. Our study opens new avenues for dissecting the biochemistry and structural biology of HIV-1 nuclear import as well as downstream events including core uncoating and potentially integration, with unprecedented detail.

## Introduction

As a lentivirus, HIV-1 can infect non-dividing immune cells, including resting CD4+ T cells, dendritic cells, and macrophages^1, 2, 3, 4, 5, 6, 7^. Upon entering the cell, the HIV-1 capsid that encapsulates its genetic material and viral enzymes, collectively termed the HIV-1 core, traverses through the cytoplasm and enters into the nucleus, where the integration of reverse-transcribed viral DNA occurs^8^. During this stage of the viral life cycle, the HIV-1 capsid is a key orchestrator, mediating multiple steps by interacting with various host factors to facilitate reverse transcription, cytoplasmic trafficking, nuclear import, and post-import trafficking to chromosomes^9, 10^. Traversing through nuclear pore complexes (NPCs) is critical for HIV-1 infection of non-dividing cells.

The HIV-1 capsid, composed of approximately 200-250 capsid protein (CA) hexamers and 12 CA pentamers, predominantly forms a conical shape with a wide-end diameter of 60 nm and a length of 120 nm^11, 12, 13, 14^, a size that significantly exceeds the ∼45 nm inner diameter of the NPC as resolved from isolated nuclear envelopes^15^. However, recent studies suggest that HIV-1 nuclear import occurs with a nearly intact capsid lattice, with uncoating occurring in the nucleus, near sites of integration^16, 17, 18, 19, 20, 21, 22^. Supporting this idea, NPCs were shown to “dilate” within cells compared to prior isolated forms, with their inner diameters approaching 65 nm^23, 24, 25, 26, 27, 28^, sufficiently large to accommodate passage of the HIV-1 capsid.

The nuclear import of HIV-1 cores is mediated by complex interactions between the capsid and components of the NPC, which facilitate directional transit through the NPC. This process involves the interaction between capsid with so-called phenylalanine-glycine (FG)-nucleoporins (Nups), such as Nup358, Nup42, Nup54, Nup58, Nup62, Nup98, POM121, and Nup153^29, 30, 31, 32, 33, 34, 35, 36, 37, 38, 39^. Recent studies show that capsid like particles (CLPs) can penetrate *in vitro* reconstituted condensates of FG-Nups, which mimic the selective barrier of the NPC’s central channel^38, 39^. However, the complex interactions involving overlapping binding pockets and different oligomerization states of various FG-Nups, as presented by the native NPC^25, 28, 33, 34, 36, 38, 40^, suggest a much more sophisticated orchestration of NPC components for the docking and traversing of HIV-1 cores. Moreover, as cores are polymorphic, exhibiting a vast heterogeneity in size and shape, whether the shape, size, and elasticity of the structure dictates its traversing through the NPC channel remains unclear.

Beyond Nups, nuclear factors including cleavage and polyadenylation specificity factor subunit 6 (CPSF6) have been shown to facilitate HIV-1 nuclear import and intranuclear trafficking^18, 41, 42, 43, 44, 45, 46^. However, the fate of the HIV-1 core post-nuclear entry remains poorly understood. Despite previous efforts to characterize HIV-1 nuclear import both in cells and *in vitro*^16, 21, 38, 39, 45, 47, 48, 49^, the low occurrence of this critical process has prevented a mechanistic understanding of the complex interplay between the core and NPC components. To overcome these challenges, we have reconstituted a functional HIV-1 nuclear import system using permeabilized CD4+ T cells and isolated viral cores, which drastically increased the occurrence of the nuclear import event. By combining targeted cryo-focused ion beam (cryo-FIB) milling with imaging techniques including cryo-correlative light and electron microscopy (cryo-CLEM) and cryo-electron tomography (cryo-ET), we effectively characterized capsid–NPC interactions and capsid integrity during nuclear import in a close-to-native environment. Our comprehensive analysis of the HIV-1 nuclear import process— from approach, docking, and traversing through the NPC to post-entry events—provides novel insights into capsid-NPC interactions and sheds light on key determinants of HIV-1 nuclear import. By analyzing close to 2000 cores, we demonstrate that successful nuclear import requires both the structural elasticity of the HIV-1 core and the expansion of NPCs, some of which undergo deformation. Brittle cores were unable to enter the NPC, while the NPC selectively facilitated the import of smaller, preferentially tube-shaped cores. CPSF6-binding-deficient cores stalled inside the NPC. Upon traversing the NPC, HIV-1 cores were found to be coated with nuclear factors, likely including CPSF6, to facilitate downstream nuclear trafficking. Collectively, our work establishes a robust nuclear import system and a highly efficient correlative cryo-ET workflow, enabling a mechanistic understanding of the interplay between the HIV-1 core and the host nuclear pore throughout the nuclear import process.

## Results

### Correlative imaging of HIV-1 core nuclear import in permeabilized T cells

Nuclear import during HIV-1 infection is rare and dynamic ^16, 21, 45, 50, 51, 52, 53^, making it extremely difficult to capture sufficient events for structural investigations. To overcome this limitation, we developed a system to recapitulate functional nuclear import using permeabilized CEM cells (an immortalized CD4+ T cell line) and isolated HIV-1 cores (Fig. 1a). Specifically, we employed digitonin permeabilization of CEM cells, which selectively permeabilizes the plasma membrane while leaving the nuclear membrane intact^54^ (Fig. 1a). Upon optimization, we determined that 0.18% digitonin provided optimal permeabilization while maintaining nuclear integrity, as confirmed by the exclusion of high-molecular-weight fluorescein isothiocyanate (FITC)-labeled dextran (500 kDa), a marker for nuclear envelope integrity (Extended Data Fig. 1a)^55^. HIV-1 cores were purified from virus-like particles (VLPs) containing mNeonGreen-labeled integrase (mNeonGreen-IN) by sucrose gradient centrifugation following spin-through delipidation^56^ (Fig. 1a). VLP cores migrated as a distinct green-fluorescence band, confirmed by cryo-EM imaging (Extended Data Fig. 1b-c). While this project initiated with VLP cores due to biosafety constraints, we eventually segued to working with cores derived from near full-length genome-containing HIV-1 particles, which will be referred to as native WT cores. Interestingly, we found that VLP cores exhibited cone-shaped and tube-shaped structures at roughly 1:1 ratio, whereas native WT cores and cores within virions contained 18% and 15% tube-shaped structures, respectively (Extended Data Fig. 2d). However, the average sizes of cone-shaped cores were similar across VLP cores, native WT cores, and virion cores, as were the sizes of tube-shaped cores (Extended Data Fig. 2e-f).

**Figure 1.**
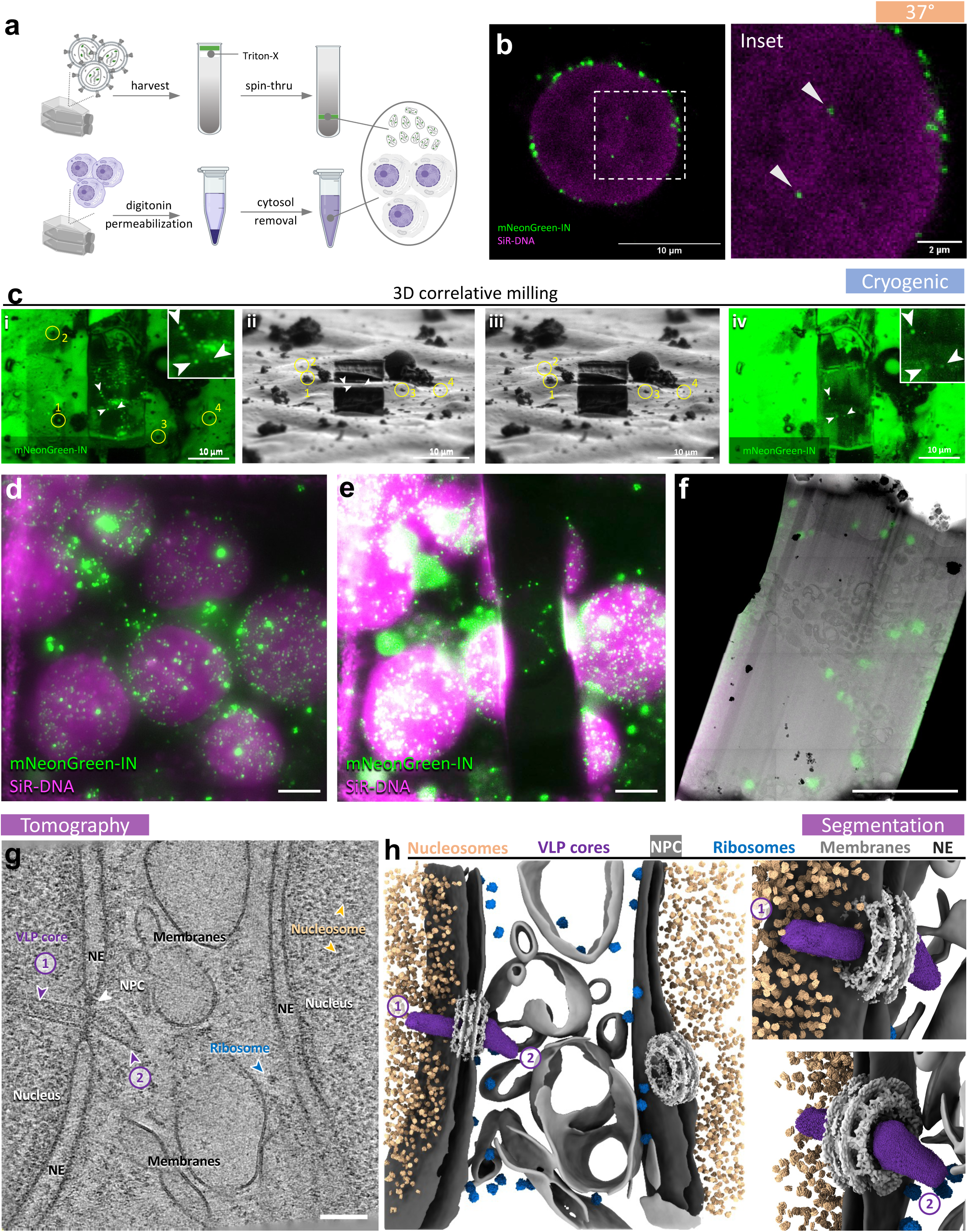
Recapitulation of functional HIV-1 nuclear import. **a,** Schematic of viral core and permeabilized T cell preparations. **b,** Confocal images of permeabilized CEM cells mixed and incubated with isolated VLP cores. VLP cores were labelled with mNeonGreen-IN (green) and the nucleus was labelled by SiR-DNA (magenta). Arrows indicate mNeonGreen-IN signals inside the nucleus. **c,** Illustration of 3D correlative cryo-FIB milling workflow. Yellow circles with numbers are ice particles used as fiducial markers for the correlation. Three white arrowheads point to targeted VLP cores. (i) Fluorescence image of the targeted nucleus after rough milling, maximum intensity projection (MIP) from 31 images, step size = 500 nm, the region of interest (ROI) is enlarged in the inset; (ii) Cryo-FIB image of the targeted nucleus after rough milling; (iii) Cryo-FIB image of the polished lamella; (iv) Fluorescence image of the polished lamella, the ROI is enlarged in the inset, MIP from 151 images, step size = 100 nm. **d-f,** Demonstration of the correlative imaging from cryo-FLM to cryo-FIB and to cryo-EM, as shown with MIP fluorescence images before (**d**) and after (**e**) milling and overlapped with the TEM overview of the same lamella (**f**). Scale bars = 5 μm. **g,** A representative tomographic slice of a correlatively-acquired tomogram of VLP core nuclear import. Two successive cores at the same NPC are indicated by purple arrowheads and numbered: No.1, an imported tube-shaped core; No.2, a cone-shaped core with wide end docked on the NPC. The NPC, ribosomes, and nucleosomes are labelled. The nucleus, nuclear envelope (NE) and membranes are annotated accordingly. Scale bar = 100 nm. **h,** The segmented volume of (**g**), shown as an overview (left) and zoomed-in views of the imported (upper right, No.1) and docked (lower right, No.2) VLP cores. VLP cores, NPCs, nucleosomes, ribosomes, NE and membranes are segmented with the indicated colours.

Intriguingly, when isolated VLP cores were mixed with permeabilized CEM cells, they were efficiently recruited to the nuclear envelope, accumulating prominently around it (Fig. 1b). Notably, the reconstituted system was fully functional for HIV-1 nuclear import, as VLP cores (green puncta) were readily detected inside the nucleus (Fig. 1b). Together, the system produced a high number of nuclear envelope-associated cores, enabling subsequent structural investigations.

Another challenge was how to precisely locate HIV-1 cores (50-100 nm) in a T cell nucleus (10 µm) to effectively characterize their nuclear import events using cryo-ET. To address this, we established a correlative workflow using cryo-fluorescence microscopy (cryo-FM), targeted cryo-FIB lamella preparation and subsequent cryo-ET imaging guided by the fluorescent signals on lamellae (Fig. 1c-f)^57, 58, 59, 60, 61, 62, 63, 64, 65, 66, 67, 68, 69, 70^. Initially, we explored the planar lift-out method, which enables the generation of wide (20–30 µm) and long (15–20 µm) lamellae from flat sample regions (Extended Data Fig. 1d)^57, 60, 62, 63, 65, 66, 68, 70^. However, to improve throughput, we adopted an automated cryo-FIB milling approach, which efficiently produces thin (100–150 nm), narrow (5–10 µm) lamellae (Fig. 1c)^58, 59, 61, 64, 67, 69^. Cryo-FIB milling was guided by core-associated mNeonGreen-IN fluorescence and SiR-DNA fluorescence marking the nucleus, to specifically target cores associated with the nuclear envelope (Fig. 1c-e). The final lamellae retained discernible mNeonGreen-IN fluorescence signal, facilitating targeted cryo-ET data collection (Fig. 1f). A representative tomogram acquired from such a correlated position on a lamella readily enumerated nuclear envelopes, NPCs, abundant ribosomes and nucleosomes (Fig. 1g-h). Notably, two successive VLP cores were seen interacting with one NPC: one tube-shaped core exiting the NPC immediately followed by a cone-shaped core docking to the same NPC (Fig. 1g-h, Supplementary video 1).

By targeting green puncta-associated positions, we were able to image HIV-1 cores at a 52% success rate (Extended Data Fig. 2, Extended Data Fig. 3a-c, and Supplementary video 2), significantly improving upon the 1–3% efficiency achieved in infected cells^21, 48^. By analyzing multiple events, we were able to discern four distinct stages of HIV-1 nuclear import: approaching, docking, traversing through the NPC, and imported (Fig. 2a-b, Extended Data Fig. 2a and Supplementary video 3). Among 467 captured VLP cores, more than 87% were associated with the T cell nucleus, either in the vicinity of an NPC or translocated into the nucleoplasm (Fig. 2c and Extended Data Fig. 2a), consistent with the robust VLP core decoration of nuclear envelopes by fluorescence light microscopy (Fig. 1b). These nuclear-associated VLP cores (total 410) were distributed across the four distinct stages: approaching (22.6%), docking (29.8%), traversing through NPCs (36.6%), and imported into the nucleus (11%) (Fig. 2c), suggesting that traversing through the NPC is a rate-limiting step, consistent with previous studies^45, 52^. Overall data statistics and experimental parameters for cryo-FIB lamella preparation and cryo-ET data collection and subtomogram averaging are detailed in Supplementary Tables 1-14.

**Figure 2.**
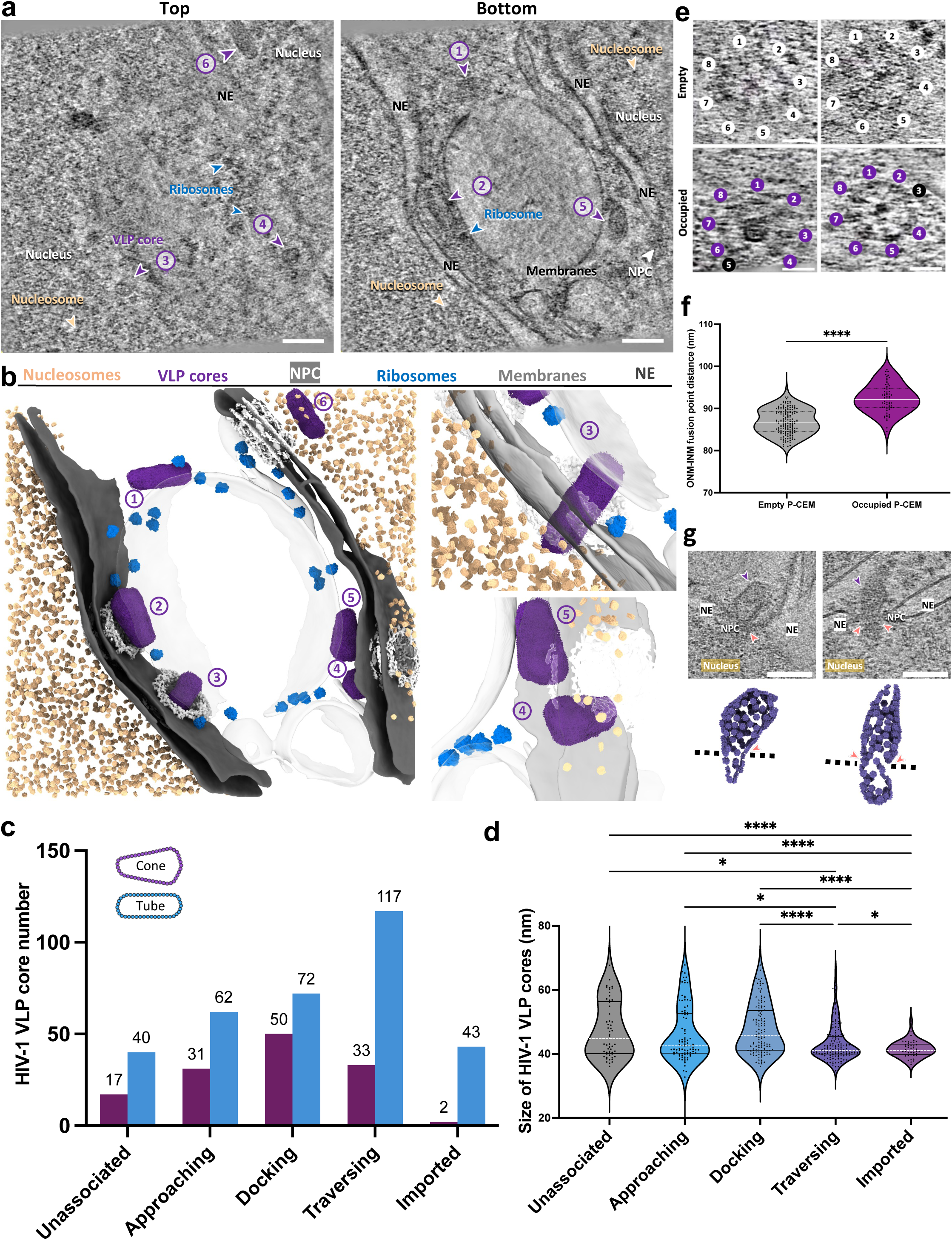
Capturing VLP cores in multiple states during nuclear import. **a,** Slices of a correlatively-acquired tomogram at the top (left) and bottom (right) of the volume featuring six VLP cores in multiple states. Cores are indicated by purple arrowheads and numbered 1 through 6. The NPC, ribosomes and nucleosomes are labelled. The nucleus, nuclear envelope (NE) and membranes are annotated accordingly. Scale bars = 100 nm. **b,** The segmented volume of (**a**) shown as an overview of six VLP cores in different states (left) and zoomed-in views of the traversing (upper right) and approaching (lower right) cores. The states of cores: unassociated tube-shaped core (No.1); docked cone-shaped core (No.2); traversing tube-shaped core (No.3); two approaching cone-shaped cores (No.4 and 5); and imported tube-shaped core (No.6). VLP cores, NPCs, nucleosomes, ribosomes, NE and membranes are segmented with the indicated colours as in Figure 1. **c,** A bar chart illustrating the composition of VLP core shapes in each state. Cone-shape cores are in purple and tube-shaped cores are in blue (Chi-square test for all, p < 0.0001). **d,** A violin plot of the statistical analysis on VLP core sizes (width measured at the wide end) in each state. The size of imported VLP cores measures 41.56 ± 2.572 nm (SE = 0.3834, n = 45), the traversing measures 43.28 ± 5.736 nm (SE = 0.4683, n = 150), the docking measures 47.80 ± 7.779 nm (SE = 0.7043, n = 122), the approaching measures 46.17 ± 8.453 nm (SE = 0.8765, n = 93), and the unassociated measures 47.40 ± 8.615 nm (SE = 1.141, n = 57). White lines represent the medians, black lines represent the quartiles, and black dots represent individual VLP cores (Brown-Forsythe and Welch ANOVA tests, * = p < 0.05, **** = p < 0.0001, ns = no significance). **e,** Tomographic slices depict the deformation of NPCs as VLP cores traverse through. NPCs in the top row, without VLP core occupancy, show an eight-fold symmetric structure (8 discernible subunits indicated by numbers). NPCs in the bottom row, with VLP cores inside, show both disruption of the eight-fold symmetry (seven discernible subunits with one offset subunit, numbered in black). Scale bars = 50 nm. **f,** A violin plot of the size distribution of NPCs with VLP cores (purple, 92.34 ± 3.336 nm, SE = 0.3987, n = 70) and without cores (grey, 86.86 ± 2.943 nm, SE = 0.2257, n = 170). White lines represent the medians, black lines represent the quartiles, and dots represent individual NPCs (t test, **** = p <0.0001). The size of NPCs is measured by the distance between the outer-nuclear-membrane (ONM) and inner-nuclear-membrane (INM) fusion points. **g,** Tomographic slices (top) and mapping back of CA hexamers (bottom) of two VLP cores depict the deformation of cores within NPCs. VLP cores are indicated by purple arrowheads, deformation points are indicated by red arrowheads and dashed black lines. NPCs, NE, and nuclei are annotated accordingly. Scale bars = 100 nm.

Closer inspection revealed that most VLP cores undergoing productive nuclear import, especially those in the traversing and imported states, were tube-shaped. Strikingly, 95% of imported VLP cores exhibited a tube-like morphology (Fig. 2c), with only two exceptions, which were comparatively narrow cone-shapes (average width: 48.6 nm) (Fig. 2c).

Furthermore, traversing and imported VLP cores were smaller than docking and approaching cores, suggesting a preference for smaller, predominantly tube-shaped cores during nuclear import (Fig. 2d). In a few instances, VLP cores appeared to disrupt the characteristic NPC 8-fold symmetry when they occupied the channel (Fig. 2e), with the majority of NPCs adopting an expanded conformation (average diameter of 92.3 nm, n = 70), compared to empty NPCs (average diameter of 86.9 nm, n = 170) (Fig. 2f). Conversely, we occasionally observed NPC-induced deformation of the capsid (Fig. 2g and Supplementary video 4-5), suggesting an interplay between the HIV-1 capsid and the NPC during nuclear import.

### Recapitulation of functional HIV-1 nuclear import using native WT cores with native nuclear pores

The selective import of tube-shaped VLP cores and the significant NPC expansion inspired us to revisit several aspects of our experimental conditions. An important question was whether NPC sizes in permeabilized cells accurately reflect those in intact cells? Another important question to address was whether the preferential import of tube-shaped cores was a mere consequence of their relative abundance in the VLP core sample compared to those from HIV-1 virions? By extension, might the presence of the viral genome inside the capsid influence nuclear import?

Given that cell permeabilization reduces cytosolic components and energy^54^, and that energy depletion has been shown to constrict NPCs^24^, we compared NPC sizes in permeabilized versus intact CEM cells. We found that the average diameter of NPCs in intact cells was 93.8 nm, significantly larger than the 86.9 nm measured in permeabilized cells (Extended Data Fig. 3d-f). This raised the possibility that the apparent preference for tube-shaped VLP cores might be biased by NPC constriction. We thus optimized our import system to a near-native condition with two advancements. First, we restored NPC size (93.4 nm, closely matching native NPCs) by supplementing permeabilized CEM cells with exogenous cytosol (rabbit reticulocyte lysate, RRL) along with an ATP-regenerating system (RRL-ATP), both of which are commonly used in nuclear import assays^71, 72^, including those for other viruses^73, 74, 75^.

Second, we used native WT cores derived from near full-length genome-containing virions (Env-defective variants of the full-length HIV-1 construct R9^76^), supplemented with mNeonGreen-IN. As mentioned above, native WT core preparations contained 18% tube-shaped cores (Extended Data Fig. 2d), consistent with previous studies^12, 13, 14^.

As demonstrated by confocal fluorescence imaging, the addition of RRL-ATP greatly increased the number of mNeonGreen-IN puncta entering the nucleus (Fig. 3a-b), reflecting enhanced nuclear import efficiency. A kinetic analysis revealed that nuclear import reached a steady-state level after 1 hour (Fig. 3b). To capture and characterize intermediate stages of nuclear import, we selected the 30-minute time point for further correlative and integrated *in situ* cryo-ET.

**Figure 3.**
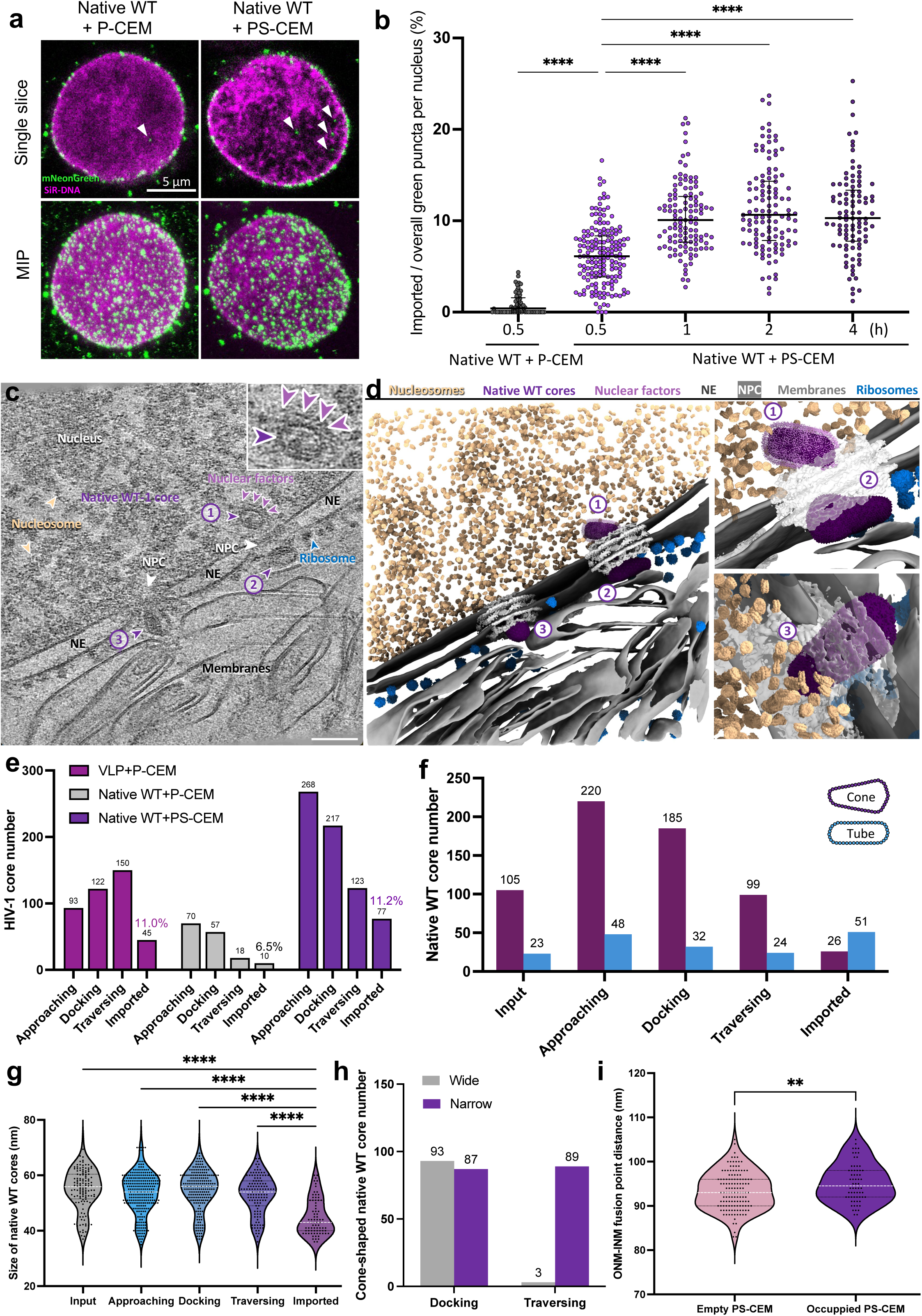
Characterisation of the nuclear import of native HIV-1 WT cores. **a,** Confocal microscopy images of permeabilized CEM cells incubated with native WT cores in the absence (P-CEM) and presence (PS-CEM) of RRL-ATP. Native WT cores are labelled with mNeonGreen-IN (green), and nuclei are labelled with SiR-DNA (magenta). Representative single Z-slice images (top) and maximum intensity projections (MIP) of Z-slices (bottom) are displayed. Arrows indicate mNeonGreen-IN signals inside the nucleus. Scale bar = 5 µm. **b,** Statistical analysis of the nuclear import of the mNeonGreen-IN puncta. The ratios represent the percentage of mNeonGreen-IN puncta localised inside the nuclei of permeabilized CEM cells under different conditions. Without RRL-ATP: 0.9% ± 1.2% (n = 61) for native WT cores. With RRL-ATP: 6.2% ± 3.3% (n = 168) for native WT cores, black lines represent medians (One-way ANOVA test and Fisher’s exact test, **** = p < 0.0001, only significant differences are shown). **c,** A representative tomographic slice of a correlatively-acquired tomogram of native WT core nuclear import. Three native WT cores are identified and indicated by purple arrowheads and numbered: No.1, an imported tube-shaped core with discernable surrounding densities (enlarged in the inset); No.2, a docked cone-shaped core with the wide end on the NPC; No.3, a cone-shaped core traversing through the NPC with the narrow end in. The NPC, ribosomes, nucleosomes, and prominent surrounding nuclear factors are labelled. The nucleus, nuclear envelope (NE), and membranes are annotated accordingly. Scale bar = 100 nm. **d,** The segmented volume of (**c**), shown as an overview (left) and zoomed-in views of the imported, docked (upper right, No.1 and No.2) and traversing (lower right, No.3) native WT cores. Native WT cores, NPCs, nucleosomes, ribosomes, nuclear factors, NE, and membranes are segmented with the indicated colours. **e,** A bar chart illustrating the distribution of HIV-1 cores in each state across three samples: VLP cores incubated with P-CEM cells (VLP + P-CEM); native WT cores incubated with P-CEM cells (native WT + P-CEM); native WT cores incubated with PS-CEM cells (native WT + PS-CEM). Imported fractions are indicated (Chi-square test for all, p < 0.0001). **f,** A bar chart illustrating the composition of native WT core shapes in each state. Cone-shaped native WT cores are in purple and tube-shaped cores are in blue (Chi-square test for all, p < 0.0001). **g,** A violin plot of the statistical analysis on the width of native WT cores (width measured at the wide end) in each state. The size of imported native WT cores measures 44.84 ± 6.519 nm (SE = 0.7429, n = 77), the traversing measures 52.97 ± 6.875 nm (SE = 0.6199, n = 123), the docking measures 54.53 ± 7.350 nm (SE = 0.4989, n = 217), the approaching measures 53.55 ± 7.432 nm (SE = 0.4540, n = 268), and the input measures 54.93 ± 7.442 nm (SE = 0.6578, n = 128). White lines represent the medians, black lines represent the quartiles, and black dots represent individual native WT cores (Brown-Forsythe and Welch ANOVA tests, **** = p < 0.0001, only significant differences are shown). **h,** A bar chart showing the orientation distribution of cone-shaped native WT cores in docking and traversing states, with the wide end in first (grey) and narrow end in first (purple) (Fisher’s exact test, p < 0.0001). **i,** A violin plot of the size distribution of PS-CEM NPCs with native WT cores (purple, 95.22 ± 4.362 nm, SE = 0.5453, n = 64) and without cores (pink, 93.39 ± 4.315 nm, SE = 0.3756, n = 132). White lines represent the medians, black lines represent the quartiles, and dots represent individual NPCs (t test, ** = p <0.01).

### Smaller native WT cores are selected for nuclear entry and recruit nuclear factors

Using this optimized system, we obtained 759 tomograms (Fig. 3c-d and Supplementary video 6) that captured 685 native WT cores at various stages of nuclear import (Fig. 3e and Extended Data Fig. 4a), enabling detailed mechanistic analyses with statistical confidence. First, consistent with our initial observations with VLP cores, smaller and tube-shaped native WT cores were selectively imported into the nucleus. Specifically, 66.2% of imported native WT cores exhibited a tubular morphology compared to 17.6% in the input population (Fig. 3f). In contrast, the ratio of cone- and tube-shaped cores remained consistent among the input and earlier stages of nuclear import. This import bias was further supported by the smaller average diameter of imported cores compared to those at prior stages of nuclear import (Fig. 3g). Notably, the variation in native WT core diameter originated from cone-shaped cores rather than tube-shaped cores, and again smaller cone-shaped cores were preferentially imported (Extended Data Fig. 4b-c). Second, while both the wide and narrow ends of cone-shaped native WT cores docked at the NPC equally well, traversing cores, however, almost exclusively penetrated the NPC via their narrow end first (Fig. 3h), in line with previous observations^77^. Third, we observed a small but significant NPC expansion (93.4 nm to 95.2 nm) upon core traversing (Fig. 3i), which, albeit less pronounced compared to permeabilized cells prior to RRL-ATP incorporation, highlighted a dynamic interplay between HIV-1 cores and the NPC.

Among the 685 captured native WT cores, 77 were located inside the nucleus. Remarkably, nearly all of these nuclear cores exhibited discernible extra densities surrounding the capsid (Fig. 3c-d, Extended Data Fig. 4a, 6a, and Supplementary video 6-11), a feature not readily observed in cores at the approaching and docking stages of nuclear import (Extended Data Fig. 4a). This observation indicated an association between the capsid and nuclear factors. To further investigate the roles of nuclear host factors in HIV-1 core import and trafficking, we performed subtomogram averaging (STA)^78, 79^ of native WT cores at distinct stages: cytoplasmic (outside), traversing, and imported. The CA hexamer structures were resolved at 12 Å, 13 Å, and 16 Å for outside, traversing, and imported native WT cores, respectively (Fig. 4 and Extended Data Fig. 5a-b), all of which aligned well with the CA hexamer model (PDB 6SKK) derived from our previous CA tubular assemblies (Fig. 4)^80^. Notably, the extents of cell-based extra densities attributable to host cell factors increased as the cores progressed inward: outside cores contained the least amount of extra density, followed by traversing cores and then imported cores, the latter of which exhibited substantial extra densities on the CA hexamer surface. This additional density, located ∼63 Å above the imported native WT core surface, coincided with CPSF6 density resolved in a cryo-EM map of the capsid-CPSF6 complex obtained using recombinant CPSF6 protein and perforated virions (Extended Data Fig. 4d-e). We further detected several cores that appeared to show signs of capsid uncoating and viral RNA/DNA release (Extended Data Fig. 6b-d, Supplementary video 8-11). The mechanism of uncoating and its molecular triggers require further investigation.

**Figure 4.**
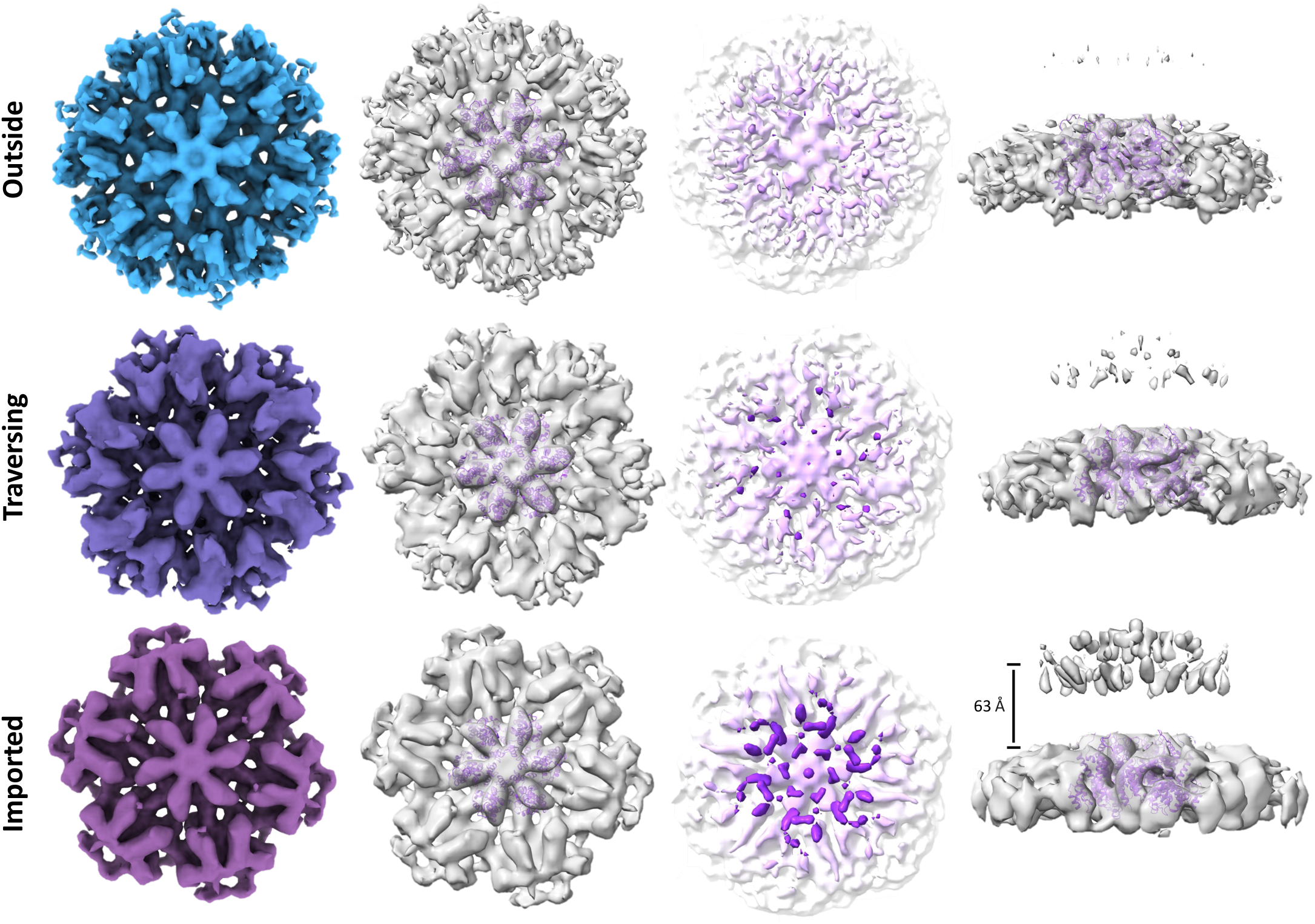
Subtomogram averaging of native WT CA hexamers during HIV-1 nuclear import. Structures of CA hexamers in capsid lattices of outside (approaching and docking combined), traversing, and imported native WT cores. Maps are aligned and contoured to the same level. The first column: top views of coloured CA hexamer density maps (contoured at 3σ); the second column: top views of CA hexamer density maps (contoured at 3σ) fitted with a CA hexamer model (PDB 6SKK); the third column: the CA hexamer density maps (contoured at 0.5σ), colored according to the height, from white (bottom) to purple (top); the fourth column: the side views of CA hexamer density maps (major body density contoured at 3σ, floating top density contoured at 0.5σ), the distance between CA hexamer and the floating top density measures approximately 63 Å for the imported native WT core.

### CPSF6-binding-deficient cores stall in the NPC

CPSF6 is thought to compete with Nup153 at the nuclear basket to release the HIV-1 core from the NPC^34, 37^. Moreover, increasing evidence suggests that CPSF6 forms condensates that co-localize with the capsid in the nucleus^43, 81, 82^, facilitating nuclear trafficking, and integration into transcriptionally-active speckle-associated domains^16, 18, 34, 35, 36, 37, 42, 43, 44, 46, 82^. We found that imported native WT cores were surrounded by extra densities consistent with recombinant CPSF6 binding to mature capsid lattices (Extended Data Fig. 4d-e). To assess how CPSF6 interactions affect HIV-1 nuclear import, we characterized native cores derived from the N74D mutant, which is deficient in CPSF6 binding^83^. Confocal fluorescence microscopy revealed a reduced nuclear import efficiency for N74D cores compared to WT cores (Fig. 5a-b), consistent with a previous immunofluorescence microscopy study of N74D-infected cells^18^. Moreover, the penetration depth of N74D cores was compromised compared to that of WT cores (Fig. 5c), in agreement with previous fluorescence imaging studies following HIV-1 infection^16, 44, 45^.

**Figure 5.**
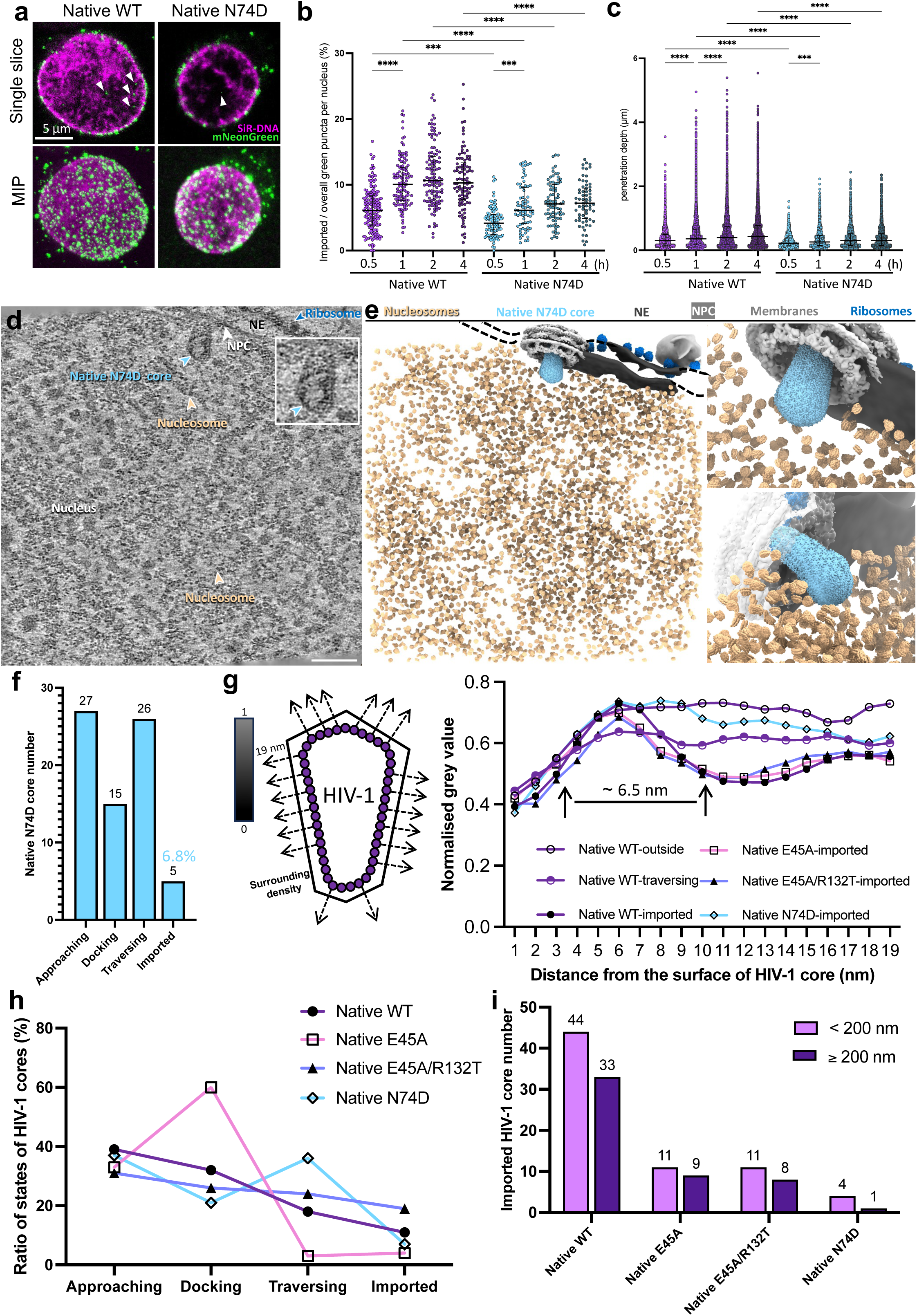
Characterisation of the nuclear import of native N74D cores. **a,** Confocal microscopy images of permeabilized CEM cells incubated with native N74D cores in presence of RRL-ATP. N74D cores are labelled with mNeonGreen-IN (green), and nuclei are labelled with SiR-DNA (magenta). Representative single Z-slice images (top) and maximum intensity projections (MIP) of Z-slices (bottom) are displayed. Arrows indicate mNeonGreen-IN signals inside the nucleus. Scale bar = 5 µm. **b,** The nuclear import efficiency of mNeonGreen-IN puncta was analysed for native WT and N74D cores. The percentage of mNeonGreen-IN puncta localized inside the nuclei of permeabilized CEM cells under different incubation times was as follows: WT 30 min: 6.2% ± 3.3% (n = 168); WT 1 h: 10.4% ± 3.7% (n = 117); WT 2 h: 11.5% ± 4.7% (n = 116); WT 4 h: 10.8% ± 4.5% (n = 98); N74D 30 min: 4.3% ± 2.2% (n = 97); N74D 1 h: 6.8% ± 3.4% (n = 72); N74D 2 h: 7.4% ± 3.1% (n = 80); N74D 4 h: 7.4% ± 3.2% (n = 66). Black lines represent medians (One-way ANOVA test and Fisher’s exact test, **** = p < 0.0001). **c,** The penetration depth of native WT and N74D cores from the nuclear envelope were enumerated as follows: WT 30 min: 0.38 ± 0.31 µm (n = 3010); WT 1 h: 0.50 ± 0.48 µm (n = 8540); WT 2 h: 0.56 ± 0.53 µm (n = 8713); WT 4 h: 0.57 ± 0.53 µm (n = 7548); N74D 30 min: 0.29 ± 0.21 µm (n = 1311); N74D 1 h: 0.35 ± 0.30 µm (n = 2257); N74D 2 h: 0.39 ± 0.32 µm (n = 2872); N74D 4 h: 0.39 ± 0.33 µm (n = 2413). Black lines represent medians (One-way ANOVA test and Fisher’s exact test, **** = p < 0.0001). **d,** A representative tomographic slice of a correlatively-acquired tomogram of native HIV-1 N74D core nuclear import. One just-imported cone-shaped native N74D core with the wide end in is identified and indicated by the light blue arrowhead. No discernable surrounding densities are observed (enlarged in the inset). The NPC, ribosomes, and nucleosomes are labelled. The nucleus, nuclear envelope (NE) and membranes are annotated accordingly. Scale bar = 100 nm. **e,** The segmented volume of (**d**), shown as an overview (left) and zoomed-in views of the just-imported native N74D core from the top (upper right) and side (lower right). The native N74D core, NPCs, nucleosomes, ribosomes, NE, and membranes are segmented with the indicated colours. **f,** A bar chart showing the distribution of native N74D cores in each state; the import fraction is annotated above the imported bar. **g,** A line chart depicting the change of density as a function of the distance from the surface of native HIV-1 cores, the first solid arrow approximately indicates the surface of the core, and the second solid arrow approximately indicates the center of the surrounding density. The cartoon on the left illustrates the measurement of grey values along the normal lines (dashed arrows) extending from the core surface, including partial densities of CA (∼3 nm) due to the resolution of images. Black density is assigned a value of 0, while white is assigned 1. Higher density corresponds to lower numerical values. For all conditions, 20 lines are drawn for each core. Numbers of HIV-1 cores analyzed: native WT-outside (n = 10), native WT-traversing (n = 10), native WT-imported (n = 10), native E45A-imported (n = 10), native E45A/R132T-imported (n = 10), and native N74D-imported (n = 5). **h,** A line chart illustrating the distribution of all native HIV-1 cores in each state in percentage (Chi-square test for all, p < 0.0001). **i**, A bar chart depicting the distribution of all imported native HIV-1 cores based on their distances from the nuclear envelope within the nucleus. The reference distance is calculated as the sum of the longest axis of the HIV-1 core (∼120 nm) and the length of the nuclear basket (∼80 nm) (Fisher’s exact test, p = 0.8641).

Corelative cryo-ET analyses revealed that native N74D cores more frequently stalled within the NPC (36%) compared to native WT cores (18%), with translated to a much-decreased import fraction (6.8%) (Fig. 5d-f, h, Extended Data Fig. 7a and Supplementary video 12), suggesting a deficiency in core release from the NPC. Furthermore, imported native N74D cores appeared comparatively “clean” with little extra surrounding densities, unlike for native WT cores. Due to the limited availability of imported N74D cores for STA, we used density profile analysis to illustrate the change in density as a function of distance from the surface of HIV-1 cores (Fig. 5g). Unlike imported native WT cores, which showed a drop in gray value starting around 6.5 nm from the capsid surface, N74D cores lacked this characteristic and showed a density profile similar to native WT cores prior to nuclear entry (Fig. 5g). Moreover, 80% of imported native N74D cores located comparatively close to the nuclear envelope (< 200 nm), whereas more than 40% of native WT cores penetrated further into the nucleus, consistent with our fluorescence data and previous studies^44, 45^ (Fig. 5i). Together, these results provide evidence that CPSF6 is, at least in part, a component of the nuclear factors bound to imported HIV-1cores, and that CPSF6 binding is essential for facilitating HIV-1 nuclear import and post-import trafficking^9, 82^.

### Capsid elasticity is essential for entering the NPC

Previous studies suggested that HIV-1 core elasticity is essential for nuclear import, and that the capsid undergoes remodeling for efficient transport through the NPC^76, 84, 85^. Brittle cores, such as those derived from the hyper stable E45A mutant, exhibit impaired nuclear import capacity, yet the nature of the interaction between these cores and the NPC remains unclear. To investigate the effect of HIV-1 core elasticity in nuclear import, we examined the nuclear import of native E45A cores and its revertant mutant E45A/R132T^76, 86, 87^. Confocal fluorescence microscopy revealed that the E45A mutation significantly decreased the number of nuclear green puncta, with an import ratio of 1.3% compared to 6.1% for native WT cores (Fig. 6a-b). As expected, the revertant mutant E45A/R132T cores, which exhibit elasticity similar to native WT cores, showed partially restored nuclear import capacity to 3.5% (Fig. 6a-b). Our observations align with previous live-cell imaging of mutant and revertant mutant nuclear import^76^, further validating our optimized nuclear import system.

**Figure 6.**
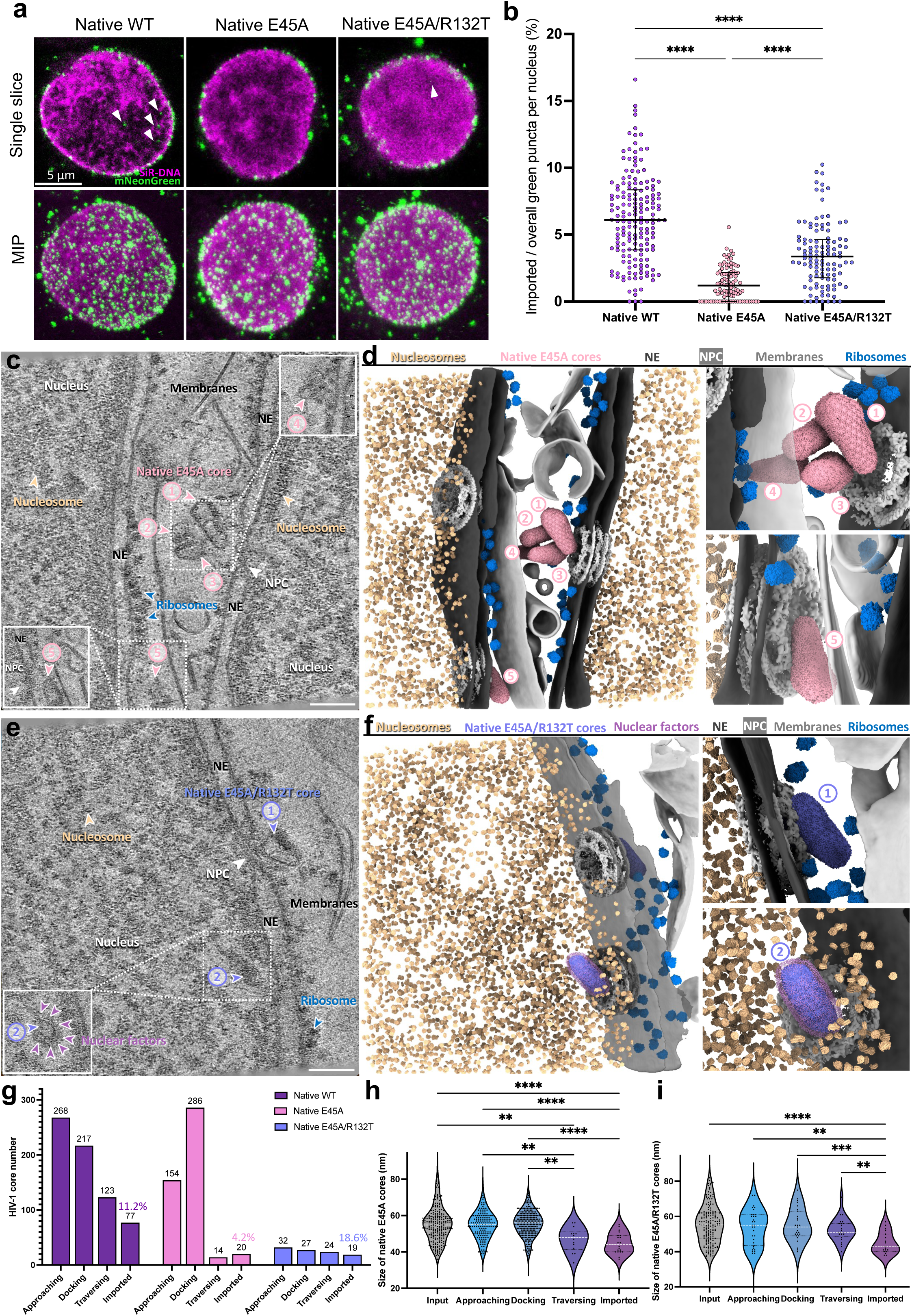
Characterisation of the nuclear import of native E45A and E45A/R132T cores. **a,** Confocal microscopy images of permeabilized CEM cells incubated with native E45A and E45A/R132T cores in presence of RRL-ATP. Cores are labelled with mNeonGreen-IN (green), and nuclei are labelled with SiR-DNA (magenta). Representative single Z-slice images (top) and maximum intensity projections (MIP) of Z-slices (bottom) are displayed. Arrows indicate mNeonGreen-IN signals inside the nucleus. Scale bar = 5 µm. **b,** Statistical analysis of the nuclear import of the mNeonGreen-IN puncta for brittle E45A revertant mutant E45A/R132T cores. The ratios represent the percentage of mNeonGreen-IN puncta localised inside the nuclei of permeabilized CEM cells under different conditions. Native WT cores: 6.2% ± 3.3% (n = 168); Native E45A cores: 1.3% ± 1.2% (n = 99); Native E45A R132T cores: 3.5% ± 2.4% (n = 111). Black lines represent medians (One-way ANOVA test, **** = p < 0.0001). **c,** A representative tomographic slice of a correlatively-acquired tomogram of native E45A cores clashing on the NPC. Five native E45A cores were identified and indicated by pink arrowheads and numbered: No.1, 3, 4, clashing cone-shaped native E45A cores; No.2, a clashing tube-shaped native E45A core; No.5, a docked cone-shaped native E45A core with the narrow end on the NPC. No. 4 and No.5 native E45A cores are shown on other slices of the same tomogram (white framed). The NPC, ribosomes, and nucleosomes are labelled. The nucleus, nuclear envelope (NE) and membranes are annotated accordingly. Scale bar = 100 nm. **d,** The segmented volume of (**c**), shown as an overview (left) and zoomed-in views of the clashed (upper right, No.1-4) and docked (lower right, No.5) native E45A cores. Native E45A cores, NPCs, nucleosomes, ribosomes, NE, and membranes are segmented with the indicated colours. **e,** A representative tomographic slice of a correlatively-acquired tomogram of native HIV-1 E45A/R132T core nuclear import. Two native E45A/R132T cores are identified and indicated by light purple arrowheads and numbered: No.1, a docked cone-shaped native E45A/R132T core with the wide end on the NPC; No.2, an imported tube-shaped native E45A/R132T core with discernable surrounding densities, also shown on another slice of the same tomogram (white framed). The NPC, ribosomes, nucleosomes, and prominent surrounding nuclear factors are labelled. The nucleus, NE and membranes are annotated accordingly. Scale bar = 100 nm. **f,** The segmented volume of (**e**), shown as an overview (left) and zoomed-in views of the docked (upper right, No.1) and imported (lower right, No.2) native E45A/R132T cores. Native E45A/R132T cores, NPCs, nucleosomes, nuclear factors, ribosomes, NE, and membranes are segmented with the indicated colours. **g,** A bar chart showing the distribution of native HIV-1 cores in each state across three samples: native WT, native E45A, and native E45A/R132T; the imported fraction in each case is annotated (Chi-square test for all, p < 0.0001). **h,** A violin plot of the statistical analysis of the size of native E45A cores (width measured at the wide end) in each state. The size of imported native E45A cores measures 44.80 ± 5.644 nm (SE = 1.262, n = 20), the traversing measures 46.79 ± 6.658 nm (SE = 1.780, n = 14), the docking measures 55.51 ± 6.993 nm (SE = 0.4135, n = 286), the approaching measures 54.82 ± 7.250 nm (SE = 0.5842, n = 154), and the input measures 55.17 ± 9.483 nm (SE = 0.6147, n = 238). White lines represent the medians, black lines represent the quartiles, and black dots represent individual native E45A cores (Brown-Forsythe and Welch ANOVA tests, ** = p < 0.01, **** = p < 0.0001, only significant differences are shown). **i,** A violin plot of the statistical analysis on the size of native E45A/R132T cores (width measured at the wide end) in each state. The size of imported native E45A/R132T cores measures 44.84 ± 5.965 nm (SE = 1.369, n = 19), the traversing measures 52.33 ± 6.670 nm (SE = 1.362, n = 24), the docking measures 55.43 ± 9.228 nm (SE = 1.776, n = 27), the approaching measures 53.88 ± 9.925 nm (SE = 1.754, n = 32), and the input measures 55.43 ± 10.53 nm (SE = 0.7763, n = 184). White lines represent the medians, black lines represent the quartiles, and black dots represent individual native E45A/R132T cores (Brown-Forsythe and Welch ANOVA tests, ** = p < 0.01, **** = p < 0.0001, only significant differences are shown).

Using correlative cryo-ET, we analyzed 474 nucleus-associated native E45A cores at different stages of nuclear translocation (Fig. 6c-d, Extended Data Fig. 8a and Supplementary video 13). Despite their morphological similarity to native WT cores (Extended Data Fig. 1c and 2d-f), their interaction with the NPC differed significantly. Tomographic analysis revealed a marked reduction not only in the number of imported native E45A cores (20/474), but surprisingly also in the number of cores traversing NPCs (Fig. 6g). As a result, there was a substantial accumulation of cores stalled at the docking stage. Interestingly, the NPC appeared to act as a bottleneck to prevent native E45A cores from entering, and on average, more native E45A cores docked to the same NPC compared to native WT cores (Extended Data Fig. 8a and c). Because native WT and E45A cores were morphologically indistinguishable (Extended Data Fig. 1c, 2e-f and 9a-b), these observations implied that the decreased elasticity of native E45A cores impeded their ability to enter the NPC channel, emphasizing the importance of capsid remodeling for efficient nuclear import. Additionally, the size of the 14 native E45A cores found inside NPCs were smaller than those in docking and approaching and had a higher percentage of tube-shaped cores (Extended Data Fig. 9c-d), but similar to those imported (Fig. 6h and Extended Data Fig. 8e), a clear distinction from native WT cores (Fig. 3g and Extended Data Fig. 9a-b). These observations reinforced that smaller-sized cores provide a selective advantage for nuclear import, particularly when the capsid lattice is inherently resistant to remodeling. As expected, native E45A/R132T cores closely mirrored the nuclear docking, traversing, and import dynamics of native WT cores (Fig. 6e-g, i, Extended Data Fig. 10, and Supplementary video 14).

### HIV-1 capsid traversing induces NPC deformation

Our analysis revealed that the NPC expands by approximately 2 nm when occupied by a native WT core (Fig. 3i). However, whether this expansion leads to structural deformation of the NPC remains unclear. We, therefore, analyzed NPC architecture from three states: native WT core-occupied NPCs (n = 43), empty NPCs (n = 95), and NPCs from uninfected intact CEM cells (n = 42) (Fig. 7a-c). Given the eight-fold symmetry of the NPC, we performed symmetry analysis to assess whether NPCs undergo deformation during native WT core traversing. Specifically, we employed template matching^78, 79^ to identify individual NPC subunits, with an NPC selection cut-off of >3 matched subunits (Extended Data Fig. 5c). The angles between adjacent template-matched subunits were measured for each NPC, with deviations exceeding 2.5° classified as deformed (Fig. 7a-c). Our analysis showed that 50% of native WT core-occupied NPCs were deformed, compared to 23% of empty NPCs and 16.6% of NPCs in native CEM cells (Fig. 7d), suggesting that the presence of cores inside the NPC promotes deformation. Further inspection of deformed empty NPCs showed they were more frequently associated with native WT cores than NPCs that remained core-unassociated (Fig. 7e). Interestingly, NPC deformation did not correlate with HIV-1 core size (Fig. 7f) nor NPC expansion (Fig. 7g). Further analysis revealed that NPC expansion upon HIV-1 core traversing was anisotropic, meaning that the structural changes were unevenly distributed, which was not due to the missing wedge effect (Fig. 7h and Supplementary video 15). In many cases, one to three subunits deviated from the rotational symmetry, contributing to NPC deformation. These observations suggested that HIV-1 core traversing not only expands the NPC but also induces structural deformation.

**Figure 7.**
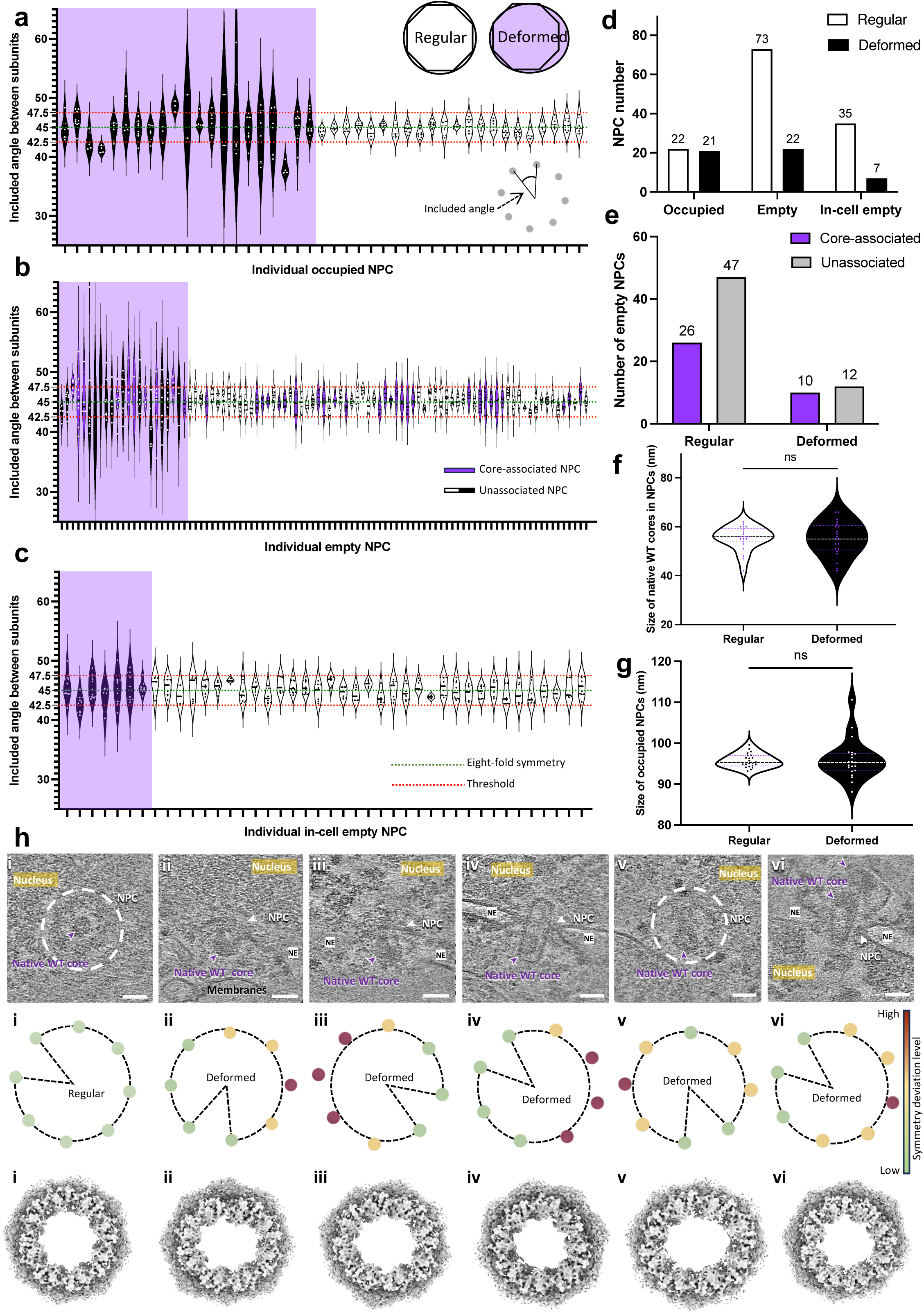
Symmetry analyses of NPCs. **a-c,** Violin plots of the included angles between adjacent subunits in PS-CEM NPCs (n = 43) occupied by native WT cores (**a**), empty PS-CEM NPCs (n = 95) (**b**), and in-cell empty CEM NPCs (n = 42) (**c**). The cartoon in the right lower of (**a**) demonstrates the measurement of included angles. The thresholds (red dashed lines) are set at 47.5° (upper) and 42.5° (lower), deviating from the eight-fold symmetry reference angle of 45° (green dashed lines). Black violin units indicate deformed NPCs while white ones indicate regular NPCs, and dots represent included angles measured in individual NPCs. Deformed NPCs, as illustrated in the cartoon in the right upper of (**a**), are grouped to the left side (purple background) of the charts for ease of comparison. For empty PS-CEM NPCs, purple violin units indicate NPCs associated with native WT cores (docking and just-imported). **d,** A bar chart depicting the distribution of NPCs in the aforementioned three conditions (Fisher’s exact test, occupied vs empty: **= p = 0.0050, occupied vs in-cell empty: ** = p = 0.0024, and empty vs in-cell empty: ns = p = 0.4983). **e,** A bar chart depicting the distribution of core-associated NPCs in regular and deformed empty NPCs (Fisher’s exact test, p = 0.4570). **f,** A violin plot of the statistical analysis on the size of native WT cores (width measured at the wide end) in regular and deformed occupied NPCs. The size of native WT cores in regular occupied NPCs measures 55.32 ± 4.854 nm (SE = 1.035, n = 22) and the size of native WT cores in deformed occupied NPCs measures 55.05 ± 6.960 (SE = 1.519, n = 21). White and black lines represent the medians, purple lines represent the quartiles, and dots represent individual native WT cores (t test, ns = no significance). **g**, A violin plot of the statistical analysis on the size of NPCs occupied by native WT cores. The regular ones measures 95.74 ± 1.684 nm (SE = 0.3591, n = 22) and the deformed measures 95.95 ± 4.841 nm (SE = 1.056, n = 21). White and black lines represent the medians, purple lines represent the quartiles, and dots represent individual occupied NPCs (t test, ns = no significance). **h**, Examples of occupied NPCs in which the eight subunits are well identified with high cross-correlation and correct orientation by template matching. The top panel depicts the six tomographic slices of native WT cores traversing through the NPC. The mid panel illustrates the symmetry analysis by rotation based on the matched coordinates of each subunit. The bottom panel shows the mapped-back model of each NPC occupied by native WT cores, using an adapted EM map EMD-51631 for better illustration. The native WT cores and NPCs are labelled (indicated by white circles in the top view in some cases). The nucleus, nuclear envelope (NE), and membranes are annotated accordingly. Scale bars = 50 nm.

## Discussion

In this work, we established a simple yet robust functional system using isolated native HIV-1 cores and permeabilized CD4+ T cells supplemented with cell lysate and an ATP regeneration system to study nuclear import in a near-native environment by correlative cryo-ET. This system not only vastly increased the abundance of HIV-1 nuclear import events compared to T cell and macrophage infection, it also enabled direct targeting of individual HIV-1 cores in the process of nuclear import, effectively capturing cores *in situ* in cryo-tomograms to an extent that was not previously feasible. As a result, an unprecedented number of HIV-1 cores were imaged at multiple stages of nuclear translocation along with their counterpart NPCs. This highly efficient system has enabled us to systematically study this dynamic process with convincing statistics. Also, the system faithfully recapitulated nuclear entry defects associated with HIV-1 mutants, demonstrating its physiological relevance.

Both confocal fluorescence imaging and cryo-ET analysis consistently demonstrate that HIV-1 cores engage robustly with the nuclear envelope, especially with NPCs, underscoring the capsid’s exceptional specificity and affinity for NPCs^38, 39^. The pronounced accumulation of cores at the nuclear envelope compared to the nucleoplasm confirms that nuclear import is a key rate-limiting step in HIV-1 infection. Our findings further reveal that HIV-1 nuclear import is highly selective, favoring the traversing of smaller cone-shaped and tube-shaped cores. This selectivity is unlikely to be an artifact of our import system, as the NPCs observed closely mirror those found in intact CEM cells (Extended Data Fig. 3f). Further evidence supporting NPCs as a selective filter comes from a previous study using resin-embedded HIV-1-infected cells, which showed that imported HIV-1 cores predominantly appeared tube-shaped or exhibited broken lattices, albeit based on a comparatively small number of observed cores^21^.

The elasticity of the HIV-1 core plays a key role in establishing infection by influencing nuclear import and subsequent uncoating^76, 77, 88^. Recent atomic force microscopy (AFM) and molecular dynamics (MD) simulation studies demonstrated that core elasticity is essential for traversing through NPCs^76, 77^. Mutations that reduce core elasticity (e.g., E45A and Q63A/Q67A), or the use of capsid inhibitors such as PF74 and Lenacapavir, appear to impair HIV-1 nuclear import^76^. However, direct visualization of the effect of capsid elasticity on nuclear import has been lacking. Here, we provide direct *in situ* evidence of HIV-1 capsid remodeling within NPCs, along with a dramatic “bottleneck” of native E45A cores at the entry of nuclear pores. Since the E45A mutation does not directly affect the capsid’s FG-binding pocket, the failure of E45A cores to enter the NPC channel reinforces the notion that structural plasticity of the capsid is critical for successful NPC traversal^84, 85^.

NPCs are known to be flexible, dynamic and capable of adapting to constricted or dilated forms depending on cell state^23, 24, 25, 26, 28, 89^. Such features facilitate the nuclear import of HIV-1 cores whose size closely matches the NPC channel. Notably, the diameters of NPCs in our import system correlate well with those in cryo-FIB milled, intact cells^23, 24, 26^, whereas isolated nuclear envelopes^15^ or permeabilized cells lacking RRL-ATP appear to constrict NPCs. We observed a small but distinct expansion (∼2 nm) of NPCs during native WT core translocation. Interestingly, when NPCs were in the constricted state due to stress, the presence of HIV-1 cores in NPCs induced a substantial expansion of nuclear pores (from 86.9 nm to 92.3 nm), suggesting that NPCs are quite flexible. In addition to the expansion, substantial NPC deformation was detected. These findings align with recent studies demonstrating that HIV-1 core traversal widens and may even crack NPCs^48, 77^. Of note, we preferred the term “deform” over “crack”.

CPSF6 has been implicated in facilitating the release of the HIV-1 cores from the NPC and downstream translocation to nuclear speckles^18, 37^. We observed native N74D cores, which are defective for CPSF6 binding, accumulated and stalled within NPCs, consistent with previously observed prolonged residence time of N74D cores at the nuclear envelope^16^. We further observed a distinct density layer surrounding imported native WT, E45A, and E45A/R132T cores, but not N74D cores. This density is consistent with the CPSF6 density observed in our cryo-EM map of capsid-CPSF6 complexes, suggesting that CPSF6 is a major component of the nuclear factors mediating core trafficking in the nucleoplasm, in line with previous cell-based assays^18, 37, 42, 43, 44, 45, 82^.

Our study uncovers unprecedented details of HIV-1 nuclear import and downstream trafficking, as summarized in Fig. 8. Our work not only advances our understanding of HIV-1 nuclear import and trafficking, it also offers a highly viable system with a robust correlative workflow applicable to investigating nuclear import mechanisms more broadly. While our study provides valuable snapshots of core traversing through the NPC and trafficking inside the nucleoplasm, future research focusing on capsid dynamics during nuclear import will be crucial. Combined with our recent cryo-ET characterization of chromatin^90^, we envision that this system may one day lead to in situ visualization of HIV-1 integration. Advanced imaging techniques such as super-resolution fluorescence microscopy, including MINFLUX, especially combined with direct capsid labeling^22, 47^, joint with the powerful correlative cryo-ET platform established herein, will offer unprecedented mechanistic insights into the dynamic process of HIV-1 infection.

**Figure 8.**
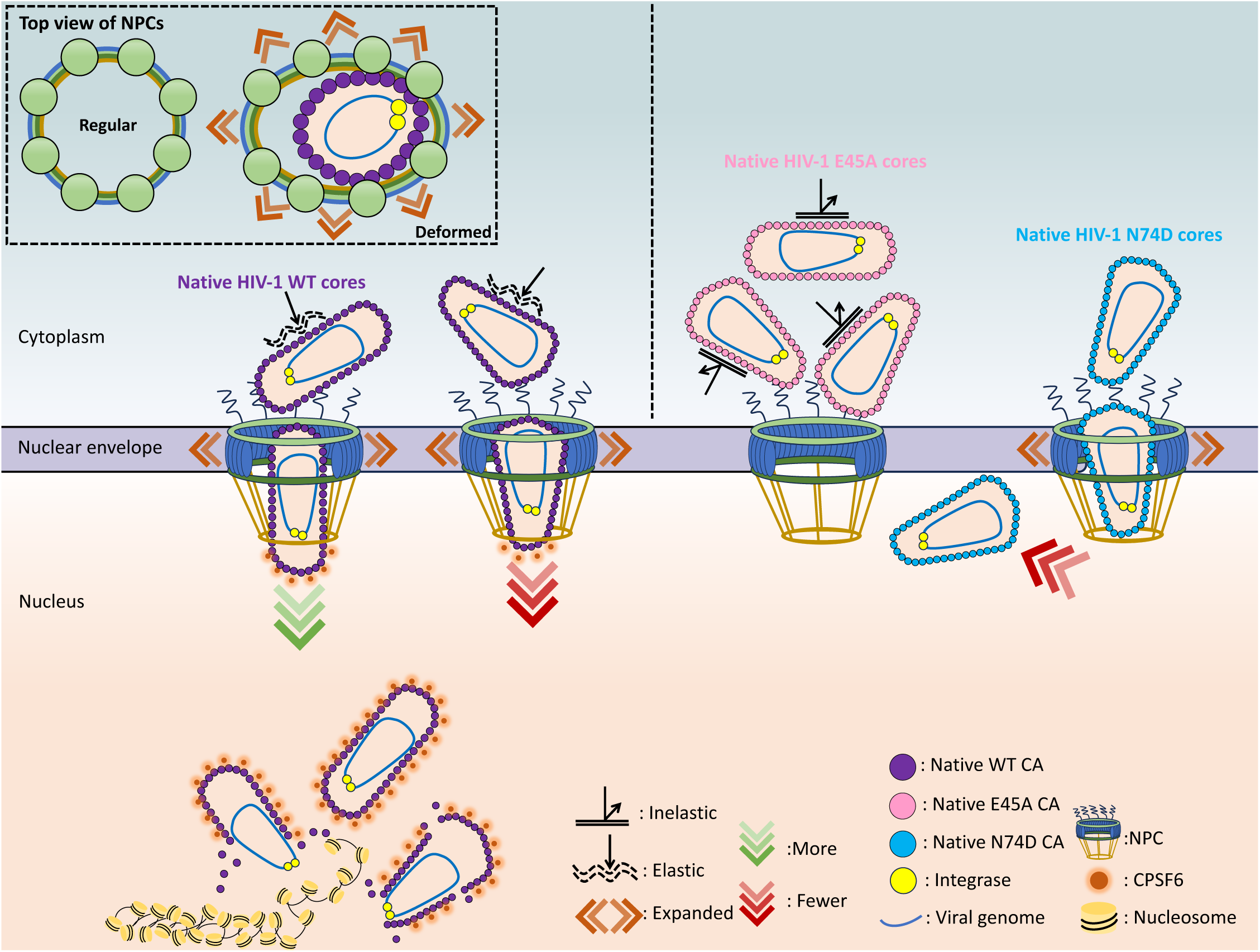
Schematic illustration of HIV-1 nuclear import. This scheme summarises key findings of our study. Native WT cores are depicted to the left: First, comparatively small and tube-shaped cores are preferentially imported through the NPC. Once inside the nucleus, imported cores are transported by nuclear factors, including CPSF6, to integration sites, where uncoating occurs, releasing the viral genome. Second, the docking orientation of HIV-1 cores is random; however, traversal through the NPC favours entry via the narrow end of cone-shaped cores. Third, NPCs expand during HIV-1 core traversal, and this process can deform the NPC, as depicted in the top left panel (framed by dashed lines). The right panel highlights the influence of capsid elasticity and CPSF6 binding on HIV-1 nuclear import, exemplified by the native E45A mutant, which has a brittle capsid, and the native N74D mutant, which is defective for CPSF6 binding. Brittle capsids cause HIV-1 cores to pile up on the NPC and disruption of CPSF6 binding hinders accumulation of core-associated extraneous nuclear density and directional inward movement.

## Material and methods

### Mammalian cell culture

Cell lines were maintained in an incubator at 37°C and 5% CO_2_. Human embryonic kidney (HEK) 293T Lenti-X cells (Takara/Clontech 632180) were cultured in DMEM (Gibco) (Sigma Aldrich) supplemented with 10% fetal bovine serum (FBS), 2 mM L-glutamine (Gibco) and 1% MEM non-essential amino acids (Gibco). CD4+ T lymphocyte CEM cells (NIH HIV reagent program/ARP-117) were cultured in RPMI-1640 (Sigma Aldrich) medium supplemented with 10% FBS and 2 mM L-glutamine (Gibco).

### HIV-1 virion production

For “in virion” core analysis, HIV-1 virus particles were produced by transfecting HEK293T cells with pNL4-3 (Env-) vectors together with NL4-3 Env expression vector pIIINL4env using Lipofectamine 2000 (Invitrogen) as previously described^91^. Culture media from transfected cells were harvested at 48 h post-transfection and cleared by filtration through a 0.45 µm polyvinylidene fluoride (PVDF) filter. The virions were concentrated by ultracentrifugation through an 8% OptiPrep density gradient (Sigma-Aldrich) (100,000 x g, Sorvall AH-629 rotor) for 1 h at 4°C. The concentrated virions were further purified by ultracentrifugation (120,000 x g, Sorvall TH-660 rotor) through a 10 – 30% OptiPrep gradient for 2.5 h at 4°C. The opalescent band was harvested, diluted with PBS, and ultracentrifuged at 110,000 × g at 4°C for 2 h. Pelleted particles were resuspended in 5% sucrose/PBS solution and stored at −80°C until use.

### Production of mNeonGreen-IN labelled HIV-1 virus-like particles (VLPs)

HIV-1 VLPs were produced by transfecting Lenti-X HEK293T cells seeded in 8 T75 flasks at ∼80% confluence. Transfections were performed using GenJet II reagent (SignaGen Laboratories). A transfection mixture was prepared by combining 40 µg of psPAX2 plasmid (psPAX2 was a gift from Didier Trono (Addgene plasmid # 12260 ; http://n2t.net/addgene:12260 ; RRID:Addgene_12260)) DNA and 10 µg of VPR-mNeonGreen-IN plasmid (was a gift from Prof. Zandrea Ambrose’s lab at University of Pittsburgh) DNA in 2 mL DMEM lacking FBS. Separately, 100 µL of GenJet II was diluted in 2 mL of DMEM without FBS, added to the plasmid DNA mixture, and incubated at room temperature (RT) for 10 min. Before transfection, each flask was refreshed with 7.5 mL of DMEM supplemented with 10% FBS, 1% nonessential amino acids (neAA), and 1% L-glutamine. Then, 500 µL of the transfection mixture was added dropwise to each flask. Cells were incubated for 48 h to allow VLP production before the supernatant was harvested.

### CPSF6 binding to SLO-treated HIV-1 VLPs

Recombinant MBP-tagged CPSF6 was purified as previously described^92^. HIV-1 VLPs for CPSF6 interaction were prepared similarly to the mNeonGreen-IN-labeled HIV-1 VLPs, except that the VPR-mNeonGreen-IN plasmid was omitted during transfection. To permeabilize the membrane of HIV-1 VLPs for CPSF6 binding, Streptolysin O (SLO) was first reconstituted by adding 100 µL of STE buffer to a vial containing 25,000–50,000 U SLO, followed by gentle mixing. The reconstituted SLO solution was then added to the VLPs at a 1.5:10 (SLO:VLP, v/v) ratio, immediately followed by the addition of IP6 to a final concentration of 1 mM. Following incubation at RT for 30 min, 5 µM MBP-tagged CPSF6 was added, and the sample was incubated at RT for another 30 min before plunge freezing.

### Production of mNeonGreen-IN labelled HIV-1 particles containing near full-length HIV-1 RNA

HIV-1 particles containing the viral RNA genome were produced by transfecting Lenti-X HEK293T cells seeded in 4 T175 flasks at ∼80% confluence. Transfections were performed using polyethyleneimine (PEI; branched; MW ∼25,000; Sigma-Aldrich). A transfection mixture was prepared by combining 24 µg of Env-defective HIV-1 constructs (R9ΔE-CA-WT or R9ΔE-CA-E45A or R9ΔE-CA-E45A/R132T or R9ΔE-CA-N74D)^76^ and 6 µg of VPR-mNeonGreen-IN plasmid DNA with 120 µg of PEI in OptiMEM medium. Mixtures were incubated at RT for 20 min prior to flask addition. After 16 h, the OptiMEM medium was replaced with fresh DMEM supplemented with 10% FBS, 1% neAA, and 1% L-glutamine. Cells were incubated for 24 h before the HIV-1-containing supernatant was harvested.

### mNeonGreen-IN labelled HIV-1 VLP and native core isolation

Virus core isolation was performed using a previously described protocol^56^ with adjustments. The virus-containing supernatant was filtered using a 0.45 μm filter (Sarstedt). The filtered supernatant was added to 38.5 ml ultracentrifuge tubes (Beckman Coulter 344058) and underlaid with 5 ml 20% sucrose in 1x STE (10 mM Tris-HCl pH 7.4, 100 mM NaCl, 1 mM EDTA pH 8.0)–1 mM IP6. The VLPs were pelleted by centrifugation using the SW32Ti rotor (Beckman Coulter Optima XPN-90) for 3 h, 30k RPM at 4 °C. After centrifugation, the supernatant and sucrose cushion were removed, and the tube was placed inverted on a tissue to dry for ∼5 min. The remaining liquid was removed by cleaning the inside walls of the tube using a tissue (Kimtech) without touching the pellet. Each VLP/virion pellet was resuspended in 400 μl 1x STE supplemented with 1 mM IP6.

A sucrose gradient was prepared in a 13.2 ml ultracentrifuge tube (Beckman Coulter 344059). All buffers were supplemented with 1x STE and 1 mM IP6. The layers were added from bottom to top using 2 ml stripettes: 2 ml 85% sucrose, 1.7 ml 70% sucrose, 1.7 ml 60% sucrose, 1.7 ml 50% sucrose, 1.7 ml 40% sucrose and 1.7 ml 30% sucrose and placed at 4 °C for ∼7-8 h. Before use, 250 μl 15% sucrose supplemented with 1% triton X-100 was added on top of the gradient and a layer of 7.5% sucrose was added on top of the triton layer to provide a barrier between the VLPs/virions and the triton before centrifugation. Finally, the 800 μl concentrated VLPs/virions were added on top of the 7.5% sucrose layer and topped off with 1x STE 1 mM IP6. The sample was separated by centrifugation using a SW41Ti rotor (Beckman Coulter Optima XPN-90) for 15-17 h, 33k RPM at 4 °C.

The sample was harvested immediately after stopping the centrifuge. The mNeonGreen band corresponding to HIV-1 cores was visualized using a blue light transilluminator and collected by side puncturing the ultracentrifugation tube using a 25 G needle (BD Microlance 3). HIV-1 cores were quantified using a p24 ELISA kit. The isolated cores were either snap frozen in single-use aliquots with liquid nitrogen and stored at −80 °C or dialyzed overnight at 4 °C against 500 ml of 1x SHE buffer (10 mM HEPES-NaOH pH 7.4 + 100 mM NaCl + 1 mM EDTA pH 8.0) supplemented with 0.8 mM IP6 using a 0.1-0.5 ml 7 kDa cutoff Slide-A-Lyzer Dialysis Cassette (Thermo Fisher Scientific) before mixing with permeabilized T cells.

### T cell membrane permeabilization

T cell membrane permeabilization was performed using cells in the exponential growth phase at a concentration of ∼1 × 10⁶ cells/mL. Cells were pelleted by centrifugation at 500 × g for 5 min at 4°C, washed twice with 10 mL ice-cold PBS, and pelleted again under the same conditions. For permeabilization, the washed pellet was resuspended in lysis buffer (20 mM HEPES, pH 7.5, 25 mM KCl, 5 mM MgCl₂, 1 mM DTT, and 1X protease inhibitor) supplemented with 0.018% digitonin. The incubation was carried out at RT with rotation for 10 min, following the protocol described previously^71^. After incubation, cells were subjected to a brief centrifugation at 200 g for 1 min, and the supernatant was removed. The pellets were resuspended in fresh lysis buffer without digitonin to eliminate residual digitonin. To evaluate nuclear integrity, digitonin-permeabilized CEM cells were incubated with 0.2 mg/mL FITC-dextran (500 kDa; Sigma-Aldrich, Cat. No. 46947) in lysis buffer for 15 min at RT. Imaging was performed using a Leica TCS SP8 confocal microscope.

### Mixing mNeonGreen-IN labelled HIV-1 cores with permeabilized T cells

The nuclei of permeabilized T cells were stained with 1 μM SiR-DNA (Spirochrome) on ice for 30 min. Digitonin-permeabilized CEM cells (400,000) were incubated with VLP cores or native HIV-1 cores (WT, E45A, E45A/R132T, or N74D) at 20 µg/mL CA concentration in a 40 µL reaction containing a buffer composed of 0.2 mM IP6, 20 mM HEPES (pH 7.5), 18.8 mM KCl, 20 mM NaCl, 0.25 mM EDTA, 3.8 mM MgCl₂, 1 mM DTT, and 1X protease inhibitor. After 30 min on ice, 10 µL of a supplement mix was added to the 40 µL reaction, achieving a final concentration of 16% (v/v) RRL (Promega, cat: L4151), 0.6 mM ATP, 0.06 mM GTP, 5 mM creatine phosphate, and 10 U/µL creatine kinase. The reaction was then incubated at 37°C for 30 min. For cryo-FIB and cryo-ET, samples were fixed with 5 mM EGS (30 minutes, RT), inactivated with 50 mM Tris-HCl (pH 7.5), and kept on ice before grid preparation.

### Confocal microscopy and fluorescence image processing

For all light microscopy imaging, the Leica TCS SP8 confocal microscope equipped with a HC PL Apo x63 MotCORR Water CS2 Objective NA 1.2 and a HyD GaAsP detector, controlled with LAS X software was used (Leica). Excitation with 488 nm, 561 nm and 633 nm laser lines were used and dynamic filter settings were applied. Z-stacks were collected with a 0.3-0.5μm step size. For both imaging and Z-stacks, a 1AU pinhole and a 1024×1024 resolution was used. Images, Z-stacks, were processed using Leica Application Suite X (Leica) and FIJI ImageJ^93^, and Arivis Vision4D.

To analyze mNeonGreen-IN puncta representing HIV-1 cores associated with nuclei or located inside nuclei, confocal image stacks were processed using Arivis Vision4D software (version 4.1.2). Nuclei were segmented using the Cellpose algorithm integrated into Arivis Vision4D. The SiR-DNA channel was used to guide segmentation. The Cellpose settings were optimized with the “cyto2” model for nuclear detection. Optimized parameters to ensure accurate identification of nuclear boundaries included a mask threshold of 2.5, a mask quality threshold of 0.2, and a nuclear diameter of 10 µm. The resulting nuclear segmentations were exported as 3D objects for further analysis. For mNeonGreen-IN puncta detection, the Blob Finder tool in Arivis was applied to the mNeonGreen fluorescence channel. The following parameters were applied to optimize detection while minimizing false positives: normalization: manual; thresholding: auto threshold; blob size: 0.35 µm; probability threshold (p, threshold): ∼30% and split sensitivity: 40%. Puncta were classified as either “imported” or “decorating” based on spatial colocalization with nuclear boundaries. Puncta entirely enclosed within a nucleus were classified as “imported,” while those partially overlapping with nuclear boundaries were categorized as “decorating.” Quantitative analyses and measurements were performed using the built-in tools in Arivis Vision4D, and data were exported for statistical analysis. To ensure reliable and accurate results, counts for each individual nucleus were inspected manually.

### Plunge freezing vitrification

The samples were plunge-frozen on glow-discharged gold finder grids, R2/2 Au G300F1 (Quantifoil), or copper grids, R2/1 Cu 300 (Quantifoil), using the Leica EM GP2 automated plunge freezer (Leica). Sample preparation was conducted in the blotting chamber at 20°C with 95% humidity. The mixture of HIV-1 cores with permeabilized CEM cells were incubated with 10% glycerol for 2 min prior to plunge freezing. A mixture of 3-6 μl of HIV-1 cores and permeabilized CEM cells (∼4000 cells/μl) was added to the carbon side of the grid, while 1 μl of PBS was added to the back side of the grid. The grids were back blotted for 6 s using filter blotting paper (Whatman) and immediately plunge frozen in liquid ethane. For control groups, 3.5 μl of native HIV-1 virions, VLP cores, native cores and MBP-tagged CPSF6 bound to the perforated VLPs were added to the carbon side of the grid, blotted from the back for 3 s, and then plunge frozen in liquid ethane. For intact CEM cells, ∼3000 cells/μl were blotted for 8 s using EM GP2 automated plunge freezer (Leica).

### Correlative cryo-FIB milling

The vitrified mixture of permeabilized CEM cells with/without supplementation of RRL-ATP and HIV-1 cores was further thinned by cryo-FIB milling to prepare lamellae, guided by cryo-CLEM in two systems. Eight grids were then loaded by a robotic delivery device (Autoloader) onto a dual-beam FIB/SEM microscope, Arctis (Thermo Fisher Scientific). This microscope is equipped with a cryogenic stage cooled to −191°C, a wide-field integrated fluorescence microscope (iFLM) system with a 100x objective (NA 0.75), and a plasma multi-ion source (argon, xenon, and oxygen), with argon used as the FIB source in this study.

Before milling, an organometallic platinum layer was deposited on the grid using the GIS system (Thermo Fisher Scientific) for 50 s. The 3D correlative milling was performed using the embedded protocol in WebUI version 1.1 (Thermo Fisher Scientific), with the milling angle adjusted to 10°. For each position, a 15-μm stack composed of fluorescence (GFP) and reflection images was acquired at a step size of 500 nm after rough milling using default parameters. The positions of targeted fluorescence spots were calculated using discernible ice chunks as fiducial markers in both SEM and FIB images, guiding the placement of lamella preparation patterns. The lamellae were produced in a stepwise sequence: i) opening at 2 nA, ii) rough milling at 0.74 and 0.2 nA, and iii) polishing at 60 pA, with the final thickness of lamellae set to 140 nm. To enhance signal detection on polished lamellae, the step size was changed to 100 nm, resulting in a 15-μm stack composed of 101 images in both GFP and far-red channels.

Twenty-five grids were loaded onto a dual-beam FIB/SEM microscope, Aquilos 2 (Thermo Fisher Scientific), equipped with a cryo-transfer system and rotatable cryo-stage cooled to - 191°C by an open nitrogen circuit. The Aquilos 2 FIB/SEM microscope was modified to accommodate a FLM system, METEOR, with a 50x objective (NA 0.8) (Delmic Cryo BV). Grids were mounted on a METEOR shuttle with a pre-tilt of 26°, followed by coating with an organometallic platinum layer using the GIS system (Thermo Fisher Scientific) for 30 s. The milling angle was set to 10°. Cells seeded in optimal positions (near the centre of the grid square) were selected for lamella production. Before milling, fluorescence stacks were collected for all selected positions in both GFP and far-red channels at a step size of 200 nm, ranging ±6 μm from the focal point. The stacks were further processed in ImageJ^94^ to enhance the signal-to-noise ratio (SNR). FIB and SEM images were then acquired and correlated with the fluorescence images using the open-source software 3D Correlation Toolbox^61^, with discernible ice chunks used as fiducial markers. Lamella preparation patterns were then placed based on the correlated positions, followed by sequential milling conducted by the software AutoTEM 5 (Thermo Fisher Scientific) from 0.5 nA (rough milling) to 0.3 nA (medium milling), 0.1 nA (fine milling), 60 pA (first polishing), and 30 pA (final polishing), with the final thickness of lamellae set to 120 nm. After final polishing, fluorescence stacks of lamellae were acquired in both GFP and far-red channels at a step size of 100 nm, ranging ±2 μm from the focal point. The light intensity and exposure time were set to 400 mW and 300 ms for the METEOR system. In total, 35 lamellae and 85 lamellae with discernible fluorescence were produced in Arctis and Aquilos 2, respectively.

For intact CEM cells, blind milling was conducted. Thinning was performed using a dual-beam FIB/SEM microscope Aquilos 2 (Thermo Fisher Scientific) equipped with a cryo-transfer system and a rotatable cryo-stage maintained at −191 °C via an open nitrogen circuit. Prior to milling, grids were mounted onto a shuttle, transferred to the cryo-stage, and coated with an organometallic platinum layer using the GIS system (Thermo Fisher Scientific) for 5–6 s. Cells located near the centres of grid squares were selected for thinning. The process was carried out stepwise using the automated milling software AutoTEM 5 (Thermo Fisher Scientific), with currents decreasing incrementally from 0.5 nA to 30 pA at 30 kV. The final thickness of the lamellae was set to 120 nm.

### Correlative planar lift-out

To perform the lift-out, a rectangular 400 × 100 mesh TEM support grid (Agar Scientific) was loaded as the receiving grid, along with the sample grid, on a 35° Aquilos 2 AutoGrid (AG) shuttle (Thermo Fisher Scientific). Before milling the bulk sample, a copper companion block (10 μm x 8 μm x 5 μm) was created from a grid bar of the receiving grid and attached to the EasyLift needle (Thermo Fisher Scientific) using FIB redeposition methods. A fluorescence overview was acquired using the iFLM system (Thermo Fisher Scientific) with a 20x objective in both GFP and far-red channels. The intensity was set to 20% and the exposure time to 200 ms. Feasible positions, or regions of interest (ROIs), with sufficient fluorescence were selected for the bulk milling.

Planar lift-out of each ROI included four main steps. These were bulk milling to create a chunk from the ROI, lift-out of the chunk, attachment of the chunk to the receiver grid, and cleaning of the front edge of the chunk (the edge that would be perpendicular to the FIB during milling to target thicknesses). A fifth step with the GIS was performed after all chunks were attached to the receiver grid. A 30 kV FIB was used in all steps. To achieve near parallel FIB milling angles relative to the surface of the bulk sample, and allow for feasible milling angles of a chunk on the receiver grid, the following formula was used to determine adequate stage tilt angles during the lift-out and attachment steps:

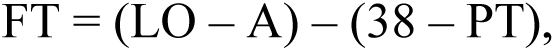

where FT is the stage tilt angle for FIB milling of a chunk, LO is the stage tilt angle during lift-out, A is the stage tilt angle during attachment to the receiver grid, 38 is the angle between the stage and FIB, and PT is the shuttle pre-tilt angle. In this planar lift-out case, FT was the stage tilt angle at −70° stage rotation (during milling of the chunks). LO and A were the stage tilt angles when the stage rotation was 110° (during the lift-out and attachment steps).

To perform the bulk milling, the stage was readjusted with a rotational angle of 110° and a stage tilt of 17° (at this position, the sample surface was perpendicular to the FIB). Bulk milling was then performed around each ROI using FIB milling currents from 1-7 nA, producing a fluorescence chunk (55 μm x 20 μm x 4 μm). High currents were used for milling thick regions, such as near grid bars, and lower currents were used for thinner regions and cleaning the edges of an ROI. Material was removed from the top, bottom, and sides of each ROI with tabs left preserved until the lift-out steps.

After bulk milling, the chunk was prepared for lift-out with the companion block on the EasyLift. The FIB was used to prepare a clean landing area on the chunk and on the block, which ensured maximum surface area contact. Once the block was contacting the chunk, attachment was achieved with 50 pA FIB current using 0.3 μm x 0.6 μm cleaning cross-section (CCS) patterns with 800-1000 ms dwell times and 2 passes. Each pattern was run for 10 s. Patterns were positioned side-by-side in groups of three and arranged along the edge of the companion block where the milling direction was from the sample towards the block. The beginning of each pattern was positioned at the point of contact between the block and chunk. After attachment, the tabs were milled, and the chunk was lifted from the bulk sample.

To perform attachment of the chunk to the receiver grid, the stage tilt was set to 9°. The FIB was used to create a landing area on the parallel grid bars and clean the edges of the chunk. The chunk was positioned between the bars and attached on the left and right sides. Attachments between the chunk and the receiver grid bars was achieved using 300 pA FIB current with 3 μm x 1 μm x 3 μm CCS patterns. The milling direction of the patterns was set towards the grid bar. Once the chunk was attached to the grid, the FIB was used to cut the companion block free of the chunk. To re-enforce the attachment of the chunk to the grid, the stage tilt was set to 17° and more attachment points were created.

With the stage tilt set to 17°, the chunk was ready for the fourth step, which was cleaning the front edge that would face the FIB during milling. After each chunk was cleaned and ready for FIB milling, the fifth and final step of the lift-out was to rotate the stage to −70°, and tilt to 5°. At this position, a 90 s GIS coating was used to condense a protective layer on the cleaned front edges of all the attached chunks on the receiver grid at the same time.

With the stage at −70° rotation and 5° stage tilt (8° milling angle with the 35° pre-tilt shuttle), milling then commenced, following the stepwise protocol described in the previous cryo-FIB milling section, guided by the remaining fluorescence. In total, five lamellae with discernible fluorescence were produced from the planar lift-out in Aquilos 2.

### Cryo-electron tomography data collection and data processing

For the experimental group, lamellae were transferred to three FEI Titan Krios G3 (Thermo Fisher Scientific) electron microscopes operated at 300 kV and equipped with a Falcon 4i detector and a Selectris X energy filter (Thermo Fisher Scientific). Objective apertures of 100 µm were inserted. Lamella overviews were generated by stitching images acquired at a magnification of 8100. The TEM lamella overviews were then correlated with the fluorescence lamella images in ImageJ^94^. Following that, tilt series were collected on the correlated sites and neighbouring areas without overlap. The collection was performed using Tomography 5 software (Thermo Fisher Scientific) at a magnification of 64k.

For permeabilised CEM cells incubated with HIV-1 VLP cores without supplementation, a total of 97 tilt series were collected with a nominal physical pixel size of 1.903 Å/pixel, 167 tilt series with a nominal physical pixel size of 1.94 Å/pixel, and 5 tilt series from the lift-out lamellae with a nominal physical pixel size of 2 Å/pixel. For tilt series from correlatively milled lamellae, the defocus value was set from −3 to −5 µm. The pre-tilts of lamellae were determined at ±10° or ±14°, and a dose-symmetric scheme was applied, ranging from −42° to +62° or −38° to +66° with an increment of 2°. A total of 53 projection images with 10 movie frames each were collected for each tilt series, with the dose rate set to 3 e/Å²/tilt, resulting in a total dose of 159 e/Å². For tilt series from lift-out lamellae, the defocus was set from −2 to - 3 µm, with no pre-tilt set. A dose-symmetric scheme was applied with a tilt range of ±60° from 0° with an increment of 2°. Tilt series were collected at a dose rate of 2.5 e/Å²/tilt in EER format, leading to a total dose of 152.5 e/Å².

For permeabilised CEM cells incubated with native WT cores, a total of 138 tilt series were collected with a nominal physical pixel size of 1.94 Å/pixel. The pre-tilts of lamellae were determined at ±10°, and a dose-symmetric scheme was applied, ranging from −44° to +64° with an increment of 2°. A total of 55 projection images with 10 movie frames each were collected for each tilt series, with the dose rate set to 2.5 e/Å²/tilt, resulting in a total dose of 137.5 e/Å², the defocus value was set from −3 to −5 µm.

For permeabilised CEM cells incubated with native WT cores supplemented with RRL-ATP, a total of 460 tilt series were collected with a nominal physical pixel size of 1.903 Å/pixel, 161 tilt series were collected with a nominal physical pixel size of 1.94 Å/pixel. The pre-tilts of lamellae were determined at ±10°/±12°, and a dose-symmetric scheme was applied, ranging from −42°/−45° to +66°+65° with an increment of 2°. A total of 53/55 projection images with 10 movie frames each were collected for each tilt series, with the dose rate set to 2.5 e/Å²/tilt, resulting in a total dose of 132.5/137.5 e/Å², the defocus value was set from −3 to −5 µm.

For permeabilised CEM cells incubated with native E45A cores supplemented with RRL-ATP, a total of 207 tilt series were collected with a nominal physical pixel size of 1.903 Å/pixel. 115 tilt series were collected with a nominal physical pixel size of 1.94 Å/pixel. The pre-tilts of lamellae were determined at ±10°, and a dose-symmetric scheme was applied, ranging from −44° to +64° with an increment of 2°. A total of 55 projection images with 10 movie frames each were collected for each tilt series, with the dose rate set to 2.5 e/Å²/tilt, resulting in a total dose of 137.5 e/Å², the defocus value was set from −3 to −5 µm.

For permeabilised CEM cells incubated with native E45A/R132T cores supplemented with RRL-ATP, a total of 122 tilt series were collected with a nominal physical pixel size of 1.903 Å/pixel. The pre-tilts of lamellae were determined at ±10°, and a dose-symmetric scheme was applied, ranging from −44° to +64° with an increment of 2°. A total of 55 projection images with 10 movie frames each were collected for each tilt series, with the dose rate set to 2.5 e/Å²/tilt, resulting in a total dose of 137.5 e/Å², the defocus value was set from −3 to −5 µm.

For permeabilised CEM cells incubated with native N74D cores supplemented with RRL-ATP, a total of 179 tilt series were collected with a nominal physical pixel size of 1.94 Å/pixel. The pre-tilts of lamellae were determined at ±10°, and a dose-symmetric scheme was applied, ranging from −44° to +64° with an increment of 2°. A total of 55 projection images with 10 movie frames each were collected for each tilt series, with the dose rate set to 2.5 e/Å²/tilt, resulting in a total dose of 137.5 e/Å², the defocus value was set from −3 to −5 µm.

For intact CEM cells, a total of 47 tilt series were collected with a nominal physical pixel size of 2.18 Å/pixel using FEI Titan Krios G2 (Thermo Fisher Scientific) electron microscope operated at 300 kV and equipped with a Gatan BioQuantum energy filter and post-GIF K3 detector (Gatan, Pleasanton, CA). A 100 µm objective aperture was inserted. Areas that include nuclei were selected for the data acquisition. Tilt series were recorded using Tomography 5 software (Thermo Fisher Scientific). The pre-tilts of lamellae were determined at ±12°, and a dose-symmetric scheme was applied, ranging from −42° to +66° with an increment of 3°. A total of 37 projection images with 10 movie frames each were collected for each tilt series, with the dose rate set to 3 e/Å²/tilt, resulting in a total dose of 111 e/Å², the defocus value was set from −3 to −5 µm.

For the control groups, grids were loaded onto a FEI Titan Krios G2 (Thermo Fisher Scientific) electron microscope operated at 300 kV and equipped with a Gatan BioQuantum energy filter and post-GIF K3 detector (Gatan, Pleasanton, CA). A 100 µm objective aperture was inserted. For the grid of near-full-genome virions and CPSF6-bound perforated VLPs, 58/100 tilt series were collected respectively using Tomography 5 software (Thermo Fisher Scientific) at a magnification of 81k with a nominal pixel size of 1.34 Å/pixel. A dose-symmetric scheme was applied with a tilt range of ±60° from 0° with an increment of 3°. Defocus was set from −1.5 to −3 µm, and the dose rate was set to 3 e/Å²/tilt. Forty-one projection images with 10 movie frames each were collected for each tilt series, resulting in a total dose of 123 e/Å². In parallel, micrographs of VLP cores were collected using EPU (Thermo Fisher Scientific) at a magnification of 105k with a nominal pixel size of 0.831 Å/pixel. A total of 59 frames were collected for each micrograph with a total dose of 22 e/Å², and the defocus was set from −3.0 to −4.0 µm. Super-resolution mode was employed, and 2000 micrographs were collected. Micrographs of HIV-1 native cores (3000 micrographs for wild-type (WT), 2000 for E45A, 2500 for E45A/R132T and 2500 for N74D) were acquired using EPU (Thermo Fisher Scientific) on a Glacios microscope. Data were collected at a magnification of 73,000x with a nominal pixel size of 2 Å/pixel. For each micrograph, a total of 40 frames were recorded with a cumulative dose of 40 e/Å². The defocus was set between - 3.0 and −4.0 µm.

### Alignment of tilt series

The frames of each tilt series were corrected for beam-induced motion using MotionCor2^95^. For the alignment and generation of tomograms, tilt series were aligned in IMOD version 4.11^96^ by patch tracking, using patches of 200 x 200 pixels and a fractional overlap of 0.45 in both X and Y. The alignment results were inspected manually, and bad frames were removed. For permeabilised CEM cells incubated with VLP HIV-1 cores without supplementation, a total of 269 tomograms were reconstructed at a binning of 6, with 97 tomograms at a pixel size of 11.418 Å/pixel, 167 tomograms at a pixel size of 11.64 Å/pixel, and 5 tomograms at a pixel size of 12 Å/pixel. For the control group of native virions, 58 tomograms were reconstructed at a pixel size of 8.04 Å/pixel. For permeabilised CEM cells incubated with native WT cores, a total of 138 tomograms were reconstructed at a binning of 6 at a pixel size of 11.64 Å/pixel. For permeabilised CEM cells incubated with native WT cores supplemented with RRL-ATP, a total of 621 tomograms were reconstructed at a binning of 6, with 460 tomograms at a pixel size of 11.418 Å/pixel, 161 tomograms at a pixel size of 11.64 Å/pixel. For permeabilised CEM cells incubated with native E45A cores supplemented with RRL-ATP, a total of 322 tomograms were reconstructed at a binning of 6, with 207 tomograms at a pixel size of 11.418 Å/pixel, and 115 tomograms at a pixel size of 11.64 Å/pixel. For permeabilised CEM cells incubated with native E45A/R132T cores supplemented with RRL-ATP, a total of 122 tomograms were reconstructed at a binning of 6 at a pixel size of 11.418 Å/pixel. For permeabilised CEM cells incubated with native N74D cores supplemented with RRL-ATP, a total of 179 tomograms were reconstructed at a binning of 6 at a pixel size of 11.64 Å/pixel. For WT HIV-1 virions, a total of 58 tomograms were reconstructed at a binning of 6 at a pixel size of 8.04 Å/pixel. For intact CEM cells, a total of 47 tomograms were reconstructed at a binning of 6 at a pixel size of 13.08 Å/pixel. SIRT-like filtering was applied to all reconstructed tomograms with 8 iterations for better visualization.

### Measurements of HIV-1 cores and NPCs

To measure the diameters of NPCs and the width of HIV-1 cores at different stages of nuclear import, tomograms were reconstructed using IMOD version 4.11.1 with a binning factor of 4. This resulted in pixel sizes of 7.612 Å/pixel, 7.76 Å/pixel, and 8 Å/pixel permeabilized CEM cells incubated with VLP cores without supplementation, 7.64 Å/pixel for permeabilized CEM cells incubated with native WT cores, 7.612 Å/pixel and 7.76 Å/pixel for permeabilized CEM cells incubated with native WT and E45A cores supplemented with RRL-ATP, 7.612 Å/pixel for permeabilized CEM cells incubated with native E45A/R132T cores supplemented with RRL-ATP, 7.76 Å/pixel for permeabilized CEM cells incubated with native N74D cores supplemented with RRL-ATP, 8.72 Å/pixel for intact CEM cells, and 5.36 Å/pixel for HIV-1 cores in native virions. The central slices of each NPC and HIV-1 core were then imported into ImageJ^94^ for precise measurements. Lines (5 lines on average) were drawn between the measuring points (ONM-INM fusion points for NPCs and the widest part for cores), and gray values were calculated along the lines. The average distance between two dropping points was recorded. Grey value measurements were taken to assess the change in density as a function of distance from the surface of native HIV-1 cores. For each core, 20 lines were drawn starting from the capsid surface. Black density was assigned a value of 0, and white was assigned a value of 1, with higher densities corresponding to lower values. The number of cores used for the measurement was as follows: native WT-outside (n = 10), native WT-traversing (n = 10), native WT-imported (n = 10), native E45A-imported (n = 10), native E45A/R132T (n = 10), and native N74D-imported (n = 5).

For the experimental groups, 467 VLP cores, 685 native WT cores, and 474 native E45A cores, 102 native E45A/R132T cores, 73 native N74D cores were measured, with the remainder being unmeasurable because they partially resided within the tomogram. For native virions, 214 cores were measured. For input cores, 131 VLP cores, 128 native WT cores, 239 native E45A cores, 184 native E45A/R132T cores, and 196 native N74D were measured directly on the motion-corrected micrographs. For NPCs with clearly definable double nuclear envelope boundaries, 240 NPCs were measured in permeabilized CEM cells without supplementation, while 196 NPCs were measured in permeabilized CEM cells supplemented with RRL-ATP, and 50 NPCs were measured in intact CEM cells.

### Template matching

To localize individual CA hexamers in the tomogram, template matching was performed using emClarity version 1.5.0.2^78^ with non-CTF-corrected tomograms binned at 6x. The procedure utilized a template derived from EMD-12452^14^, which was low-pass filtered to 30 Å. Exhausting search was applied by giving a peak threshold of 2,000 in the small cropped-out tomogram region (150 nm^3^) containing HIV-1 cores. HIV-1 CA hexamer peaks were selected with the MagpiEM tool (available at https://github.com/fnight128/MagpiEM), and particle selection was identified based on the geometric constraints of the capsid lattice. Any hexamers that did not conform to the expected hexagonal lattice geometry were automatically excluded, followed by manual inspection to ensure the selection precision (see Extended Data Fig. 5a). To localize 80S ribosomes and nucleosomes, the same procedure was used without the inspection by MagpiEM, employing low-pass-filtered maps derived from EMD-1780^97^ and EMD-16978^90^, respectively. To match the subunits of NPCs, 1/8 of the original density map EMD-11967^21^ was cropped out based on the 8-fold symmetry and further low-pass filtered to 60 Å. The same exhausting search was employed by giving a peak threshold of 200 in the small cropped-out tomogram region (300 nm^3^) containing the NPC. NPC subunit peaks were initially examined in Chimera by thresholding the cross-correlation value and then selected with the MagpiEM tool. Particle selection was identified based on the geometric constraints of NPC symmetry. Any subunit that did not align with the expected symmetry of the NPC and orientation were automatically excluded, followed by manual inspection with original tomograms to ensure selection precision (see Extended Data Fig. 5c).

### Subtomogram averaging

The 3D alignment and averaging of hexameric CA were refined through progressive binning steps from 6 to 2, employing emClarity version 1.5.3.10^79^ and maintaining C6 symmetry throughout the alignment process. The final density map at 2x binning was enhanced by sharpening with a B-factor of −10. For the CA hexamer of outside (approaching and docking) native WT cores, 128 tomograms were selected, and 11,915 particles were used. For the CA hexamer of native WT cores in traversing state, 54 tomograms were selected, and 5,545 particles were used. For the CA hexamer of imported native WT cores, 51 tomograms were selected, and 7,825 particles were used. The resolution of the reconstructed structure was calculated using a gold-standard Fourier shell correlation (FSC) cut-off of 0.143. The final resolution was determined at 11 Å, 12 Å, and 16 Å for CA hexamer structures of native WT cores in outside, traversing, and imported states, respectively (Extended Data Fig. 4d). Structural fitting was conducted in ChimeraX^98^, enabling detailed comparison of the density maps with the PDB 6SKK^80^.

### Segmentation

To enhance the segmentation, reconstructed bin6 tomograms were corrected for the missing wedge and denoised using IsoNet version 0.2^99^, applying 35 iterations with sequential noise cut-off levels of 0.05, 0.1, 0.15, 0.2, and 0.25 at iterations 10, 15, 20, 25, and 30, respectively. Membranes and nuclear envelopes in all tomograms were initially segmented using MemBrain-seg^100^, then imported into ChimeraX^98^ for manual cleaning and polishing. For the top view of nuclear envelopes, segmentation was carried out in Amira (ThermoFisher Scientific).

Nucleosomes and 80S ribosomes were mapped back to the tomograms with segmented membranes using ChimeraX^98^ and ArtiaX^101^, based on their positions and orientations after template matching. NPCs were manually placed back based on the positions of holes (occupied by NPCs) on segmented nuclear envelopes. HIV-1 cores were mapped back based on the refined coordinates and orientations of CA hexamers from subtomogram averaging, with missing CA hexamers manually placed back using the in-situ CA hexamer structures resolved in this study. CA pentamers were only placed back on cores with most CA hexamers matched and the five surrounding hexamers identified, and then modelled based on EMD-13422^14^. For deformed HIV-1 cores, CA hexamers were mapped back using the in-situ CA hexamer structure of HIV-1 cores in the traversing state. For better visualization, 80S ribosomes depicted in the segmented volume were generated by applying a low-pass filter with a cut-off of 15 Å to the original model EMD-1780^97^. Nucleosomes and NPCs were depicted using EMD-16978^90^ and EMD-11967^21^, respectively. The viral RNA/DNA was traced along the prominent string-shaped densities and segmented in ChimeraX using ArtiaX.

### Symmetry analysis of NPCs

To examine the symmetry of NPCs, template matched and cleaned coordinates of individual NPCs were first visualized with MagpiEM tool (available at https://github.com/fnight128/MagpiEM). NPCs meeting the following two criteria were retained: No.1, at least three subunits were clearly matched with correct orientation; No.2, at least one diameter was definable between two opposing subunits. Then, the direct distance between two adjacent subunits (assumed as subunit 1 and subunit 2 for simplification) was measured as *D*, and the distance between subunit 1 and its opposing subunit was measured as *d_1,_* and the distance between subunit 2 and its opposing subunit was measured as *d_2,_* in cases where only one diameter could be measured, *d_1_* and *d_2_* were set to the same value. Then the two radii were calculated as *r_1_* and *r_2_,* followed by the calculation of the included angle (θ) between these two adjacent subunits by this formula, also see in the Extended Data Fig. 5b:

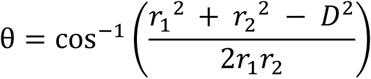

NPCs with included angles calculated larger than 47.5° or smaller than 42.5° were regarded as deformed.

### Statistical analysis

To assess the significance of size differences across HIV-1 cores in multiple states, Brown-Forsythe and Welch ANOVA tests were employed. Chi-square test was applied to determine whether the distributions of cone-shaped and tube-shaped HIV-1 cores were independent of their states (Supplementary Table 1-6). Fisher’s exact test was used to examine whether the orientation of cone-shaped HIV-1 cores was independent of the two states: docking and traversing (Supplementary Table 7-11). The significance of size differences in NPCs was assessed using an unpaired two-tailed t-test and One-way ANOVA test, including 70 HIV-1 occupied NPCs and 170 empty NPCs in permeabilized CEM cells, 64 HIV-1 occupied NPCs and 132 empty NPCs in permeabilized and supplemented CEM cells, and 50 empty NPCs in intact CEM cells. The significance of size differences in HIV-1 cores across all groups was also assessed using an unpaired two-tailed t-test and One-way ANOVA test. Fisher’s exact test and Chi-square test were used to examine the import efficiency of all HIV-1 cores, as well as to examine the distribution of all HIV-1 cores in terms of shape and states in all samples, and to assess the symmetry analysis of NPCs. Unpaired two-tailed t-test was applied to assess the difference in size between regular and deformed occupied NPCs and HIV-1 cores in them. All statistical analyses were calculated and plotted using Prism 10.

## Data availability

All data needed to evaluate the conclusions in the paper are present in the paper and/or the Supplementary information and source data are provided with this paper. The subtomogram averaged maps for the CA hexamers from native WT cores in outside, traversing, and imported states are deposited in the public database EMDB under the accession codes: EMD-52888 (outside), EMD-52887 (traversing), EMD-52889 (imported).

## Code availability

The scripts used in this study and relevant codes are deposited in GitHub [https://github.com/fnight128/MagpiEM] and Zenodo [https://doi.org/10.5281/zenodo.8362772].

## Acknowledgements

We thank Dr. Loic Carrique, Dr. Helen Duyvesteyn, Dr. David Owen, Dr. James Bancroft, Mr. Edward Drydale, and Dr. James Gilchrist for their support in data collection. We thank Dr. Yuta Hikichi and Dr. Eric O. Freed for kindly providing HIV-1 virions. We thank Mr. Frank Nightingale for the MagpiEM software. We thank Dr. Luiza Mendonça, and Dr. Joshua Hope for their suggestions in the sample optimization. Dr. Yao Shen is further supported by a CIHR fellowship from the Canadian Institutes of Health Research (funding reference number: 194032), followed by an EMBO fellowship (ALTF 96-2024). We acknowledge The Oxford Particle Imaging Centre (OPIC) for access of cryo-FIB/SEM instruments (Arctis and Aquilos 2) and cryo-EM instrument (Krios). We acknowledge Oxford Cellular Imaging Core Facility (CICF) for access of fluorescence microscopes and imaging analysis software. We acknowledge Diamond Light Source for access and support of the cryo-EM facilities at the UK national electron Bio-Imaging Centre (eBIC), (proposal NT29812). Computation was performed at the Diamond Light Source and Oxford Biomedical Research Computing (BMRC) facility supported by the Wellcome Trust Core Award Grant Number 203141/Z/16/Z with additional support from the NIHR Oxford BRC. This work was supported by US National Institutes of Health grants P50AI150481, R21AI184080 and R01AI052014, the UK Wellcome Trust Investigator Award 206422/Z/17/Z, the UK Biotechnology and Biological Sciences Research Council grant BB/S003339/1 and ERC AdG grant 101021133.

## Author contributions

P.Z. conceived the research. Z.H., Y.S., S.F., and P.Z. designed the experiments. Y.S. prepared most samples. Y.S. conducted most of fluorescence microscopy. S.F. helped in the sample preparation and fluorescence microscopy. J.S., and J.S. helped with HIV-1 core preparation. N.H. prepared CPSF6-VLPs sample. Z.H. performed the cryo-CLEM, cryo-FIB, and cryo-ET. Z.H. collected the tilt series. Z.H. and Y.S. carried out reconstruction of tomograms. Y.S. collected the micrographs of input cores. C.T. and S.N. performed the correlative lift-out and cryo-ET. Z.H. carried out the segmentation of tomograms, subtomogram averaging, and symmetry analysis. L.C. helped in the subtomogram averaging. J.X. carried out subtomogram averaging for CPSF6-VLPs. Z.H. and Y.S performed the statistical analyses. Z.H., Y.S. prepared most figures with S.F.’s help. Z.H., Y.S., S.F., and P.Z. wrote the manuscript, C.A. and A.N.E contributed to the revision, with help from all co-authors.

