## Supplementary figures and tables for "Correlative In Situ Cryo-ET Reveals Cellular and Viral Remodeling Associated with Selective HIV-1 Core Nuclear Import"

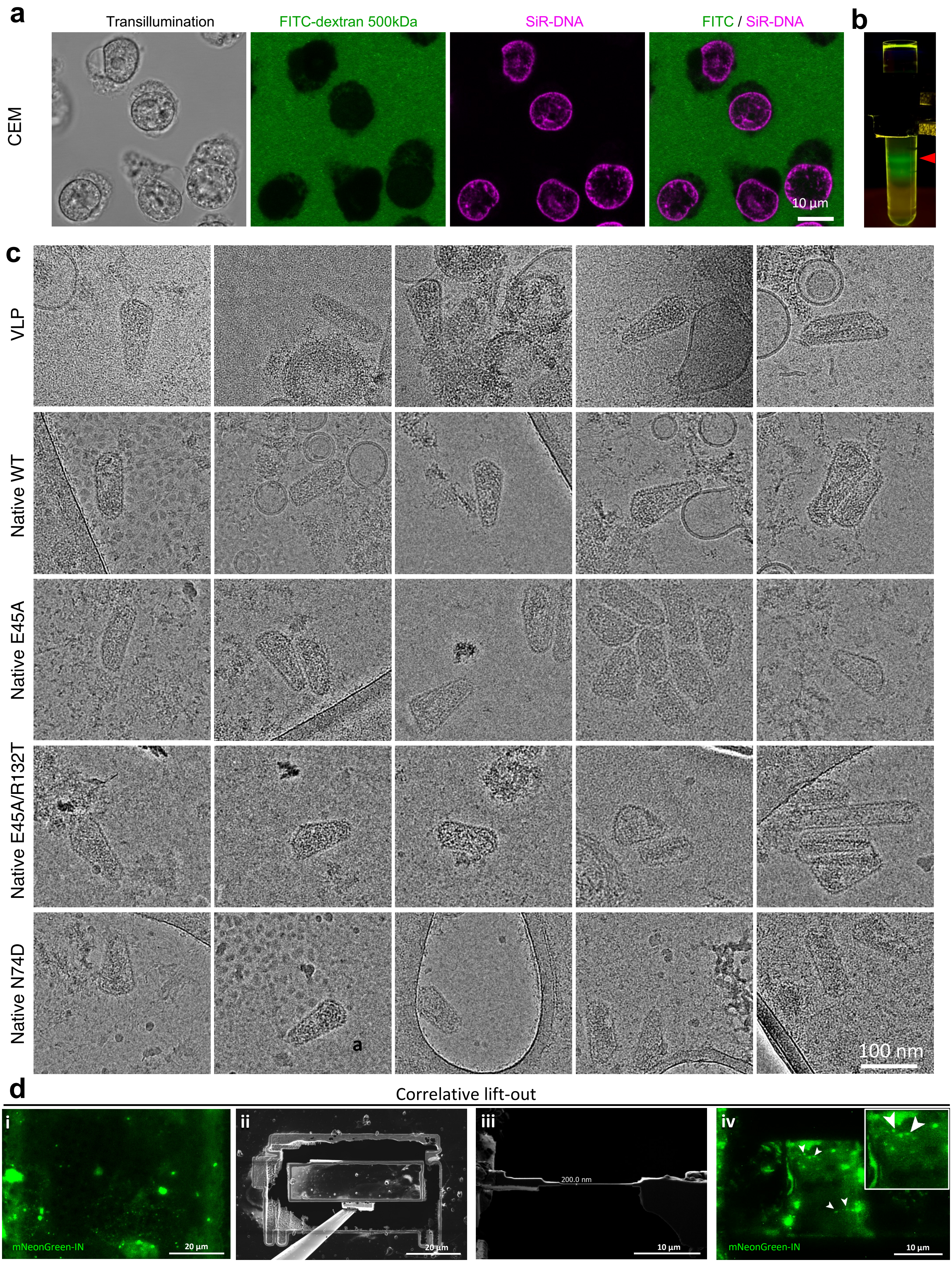
a
Transillumination
FITC-dextran 500kDa
SiR-DNA
FITC / SiR-DNA
CEM
10 µm
b
c
VLP
Native WT
Native E45A
Native E45A/R132T
Native N74D
a
100 nm
d
Correlative lift-out
i
mNeonGreen-IN
20 µm
ii
20 µm
iii
200.0 nm
10 µm
iv
mNeonGreen-IN
10 µm

**Extended Data Fig. 1 | Sample preparation and characterisation of digitonin-permeabilized CEM cells and isolated HIV-1 cores**. **a**, Representative confocal microscopy images of digitonin-permeabilized CEM cells incubated with 500 kDa FITC-dextran. Channels shown include transillumination, FITC-dextran, SiR-DNA, and a merged image of SiR-DNA with FITC-dextran. Scale bar = 10 µm. **b**, mNeonGreen-IN labeled VLP core bands after 'spin-thru' detergent treatment. The red arrowhead indicates the top band extracted for this study, while the green band below represents aggregated cores, which were discarded. **c**, Representative electron micrographs of isolated HIV-1 cores derived from VLPs and native WT, E45A, E45A/R132T, and N74D variants. Scale bar = 100 nm. **d,** Illustration of correlative planar lift-out workflow. (i) Fluorescence image of the targeted nucleus before lift-out, MIP from 20 images, step size = 500 nm; (ii) Cryo-FIB image of the lift-out slab with an EasyLift needle attached; (iii) Cryo-FIB image of the polished lamella; (iv) Fluorescence image of the polished lamella, MIP from 20 images, step size = 500 nm. White arrowheads point to the targeted HIV-1 cores, region of interest is enlarged in the inset.

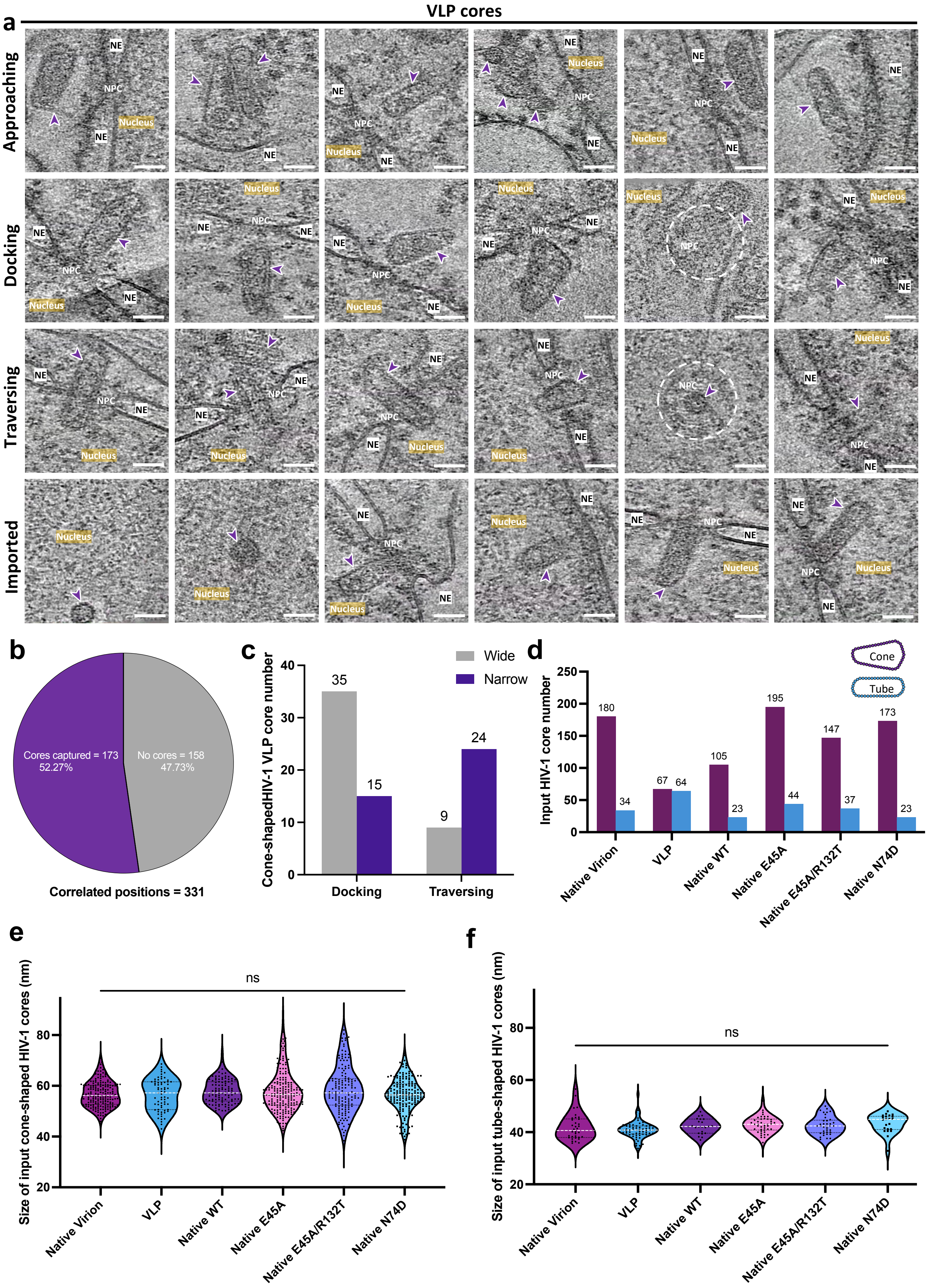

a
VLP cores
Approaching
Docking
Traversing
Imported
NE
NPC
Nucleus
b
Cores captured = 173
52.27%
No cores = 158
47.73%
Correlated positions = 331
c
Cone-shapedHIV-1 VLP core number
Wide
Narrow
35
15
9
24
Docking
Traversing
d
Input HIV-1 core number
Cone
Tube
180
34
67
64
105
23
195
44
147
37
173
23
Native Virion
VLP
Native WT
Native E45A
Native E45A/R132T
Native N74D
e
Size of input cone-shaped HIV-1 cores (nm)
ns
f
Size of input tube-shaped HIV-1 cores (nm)
ns

**Extended Data Fig. 2 | Characterisation of VLP core nuclear import**. **a**, Gallery of VLP cores in multiple states during nuclear import. Six representative tomographic slices from each state are showcased. VLP cores are indicated by purple arrowheads, the nucleus, NE, and NPC (indicated by white circles in the top view in some cases) are annotated accordingly. Scale bars = 50 nm. **b**, A pie chart illustrating the effectiveness of cryo-CLEM, showing that over 52% of correlated fluorescence positions contain VLP cores in tomograms ("cores captured"). Multiple tomograms were acquired per correlated position (e.g., at the center and surrounding areas, with a maximum of four tomograms per position). A position is classified as "cores captured" if at least one of the acquired tomograms contains a core. **c**, A bar chart showing the orientation distribution of cone-shaped VLP cores in docking and traversing states, with the wide end in first (grey) and narrow end in first (purple) (Fisher's exact test, $p = 0.0003$). **d**, A bar chart illustrating the composition of all input core shapes in each sample. Cone-shape cores are in purple and tube-shaped cores are in blue. **e**, A violin plot of the statistical analysis on the size of all input cone-shaped cores (width measured at the wide end) in each state. The size of native cone-shaped virion cores measures 56.79 $\pm$ 4.935 nm (SE = 0.3678, n = 180), the VLP measures 56.43 $\pm$ 6.199 nm (SE = 0.7573, n = 67), the native WT measure 57.71 $\pm$ 4.758 nm (SE = 0.4644, n = 105), the native E45A measures 57.89 $\pm$ 8.162 nm (SE = 0.5845, n = 195), the native E45A/R132T measures 58.66 $\pm$ 9.166 nm (SE = 0.7560, n = 147), and the native N74D measures 56.46 $\pm$ 6.644 nm (SE = 0.5052, n = 173). White lines represent the medians, black lines represent the quartiles, and black dots represent individual cone-shaped HIV-1 cores (One-way ANOVA test, ns = no significance). **f**, A violin plot of the statistical analysis on the size (width measured) of all input tube-shaped HIV-1 cores in each state. The size of tube-shaped HIV-1 virion cores measures 41.91 $\pm$ 5.056 nm (SE = 0.8672, n = 34), the VLP measures 40.95 $\pm$ 3.314 nm (SE = 0.4143, n = 64), the native WT measure 42.27 $\pm$ 2.875 nm (SE = 0.5996, n = 23), the native E45A measures 42.84 $\pm$ 3.116 nm (SE = 0.4751, n = 44), the native E45A/R132T measures 42.63 $\pm$ 3.460 nm (SE = 0.5688, n = 37), and the native N74D measures 43.40 $\pm$ 3.590 nm (SE = 0.7655, n = 23). White lines represent the medians, black lines represent the quartiles, and black dots represent individual tube-shaped HIV-1 cores (One-way ANOVA test, ns = no significance).

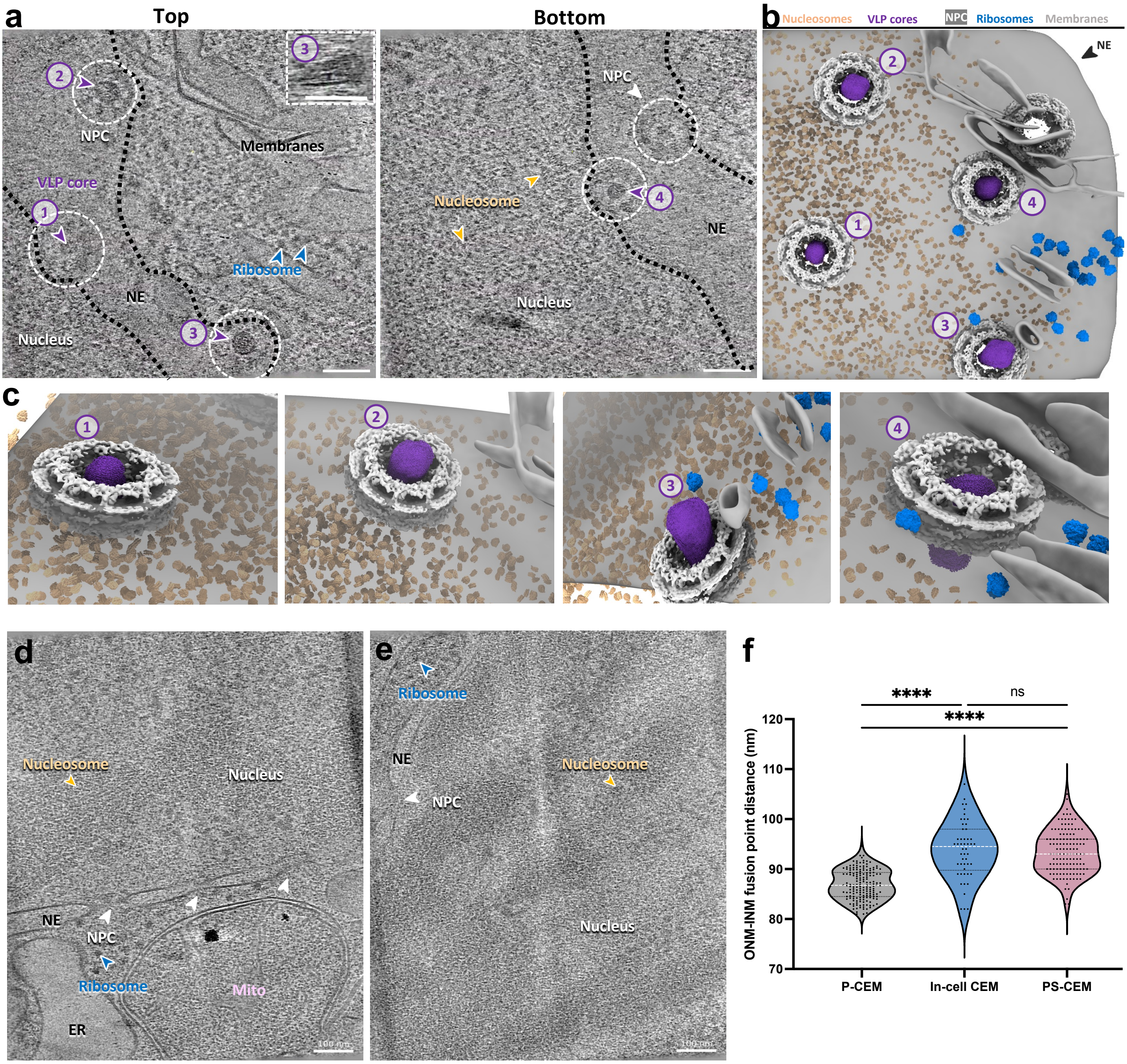


**Extended Data Fig. 3 | The interplay between HIV-1 VLP cores and NPCs**. **a**, Tomographic slices of a correlatively-acquired tomogram at the top (left) and bottom (right) of the volume showing the face-on views of NPCs occupied by VLP cores. VLP cores are indicated by purple arrowheads and numbered (1-3 on the left and 4 on the right), along with one empty NPC (white arrowhead) (right). The inset depicts the side view (XZ) of No.3 core in the NPC. NPCs are indicated by white dashed circles. The ribosomes, nucleosomes, nucleus, and nuclear envelope (NE) are annotated accordingly. The boundaries of NE are indicated by black dashed lines. Scale bars = 100 nm. **b**, **c**, The segmented volume of (**a**) shown as an overview of four VLP cores in NPCs (**b**) and zoomed-in tilted views of these four VLP cores at various depths within the NPC during traversal (**c**). VLP cores, NPCs, nucleosomes, ribosomes and NE are segmented with the indicated colours. **d**, **e**, Tomographic slices of two representative tomograms of CEM cells showing in-cell NPCs. The ribosomes, nucleosomes, nucleus, mitochondria (Mito), and NE are annotated accordingly. **f**, A violin plot of the size distribution of P-CEM, in-cell, and PS-CEM NPCs. The P-CEM measures 86.86 ± 2.943 nm (SE = 0.2257, n = 170), the in-cell measures 93.84 ± 5.744 nm (SE = 0.8123, n = 50), and the PS-CEM measures 93.39 ± 4.315 nm (SE = 0.3756, n = 132). White lines represent the medians, black lines represent the quartiles, and dots represent individual NPCs (One-way ANOVA test, **** = p <0.0001, ns = no significance).

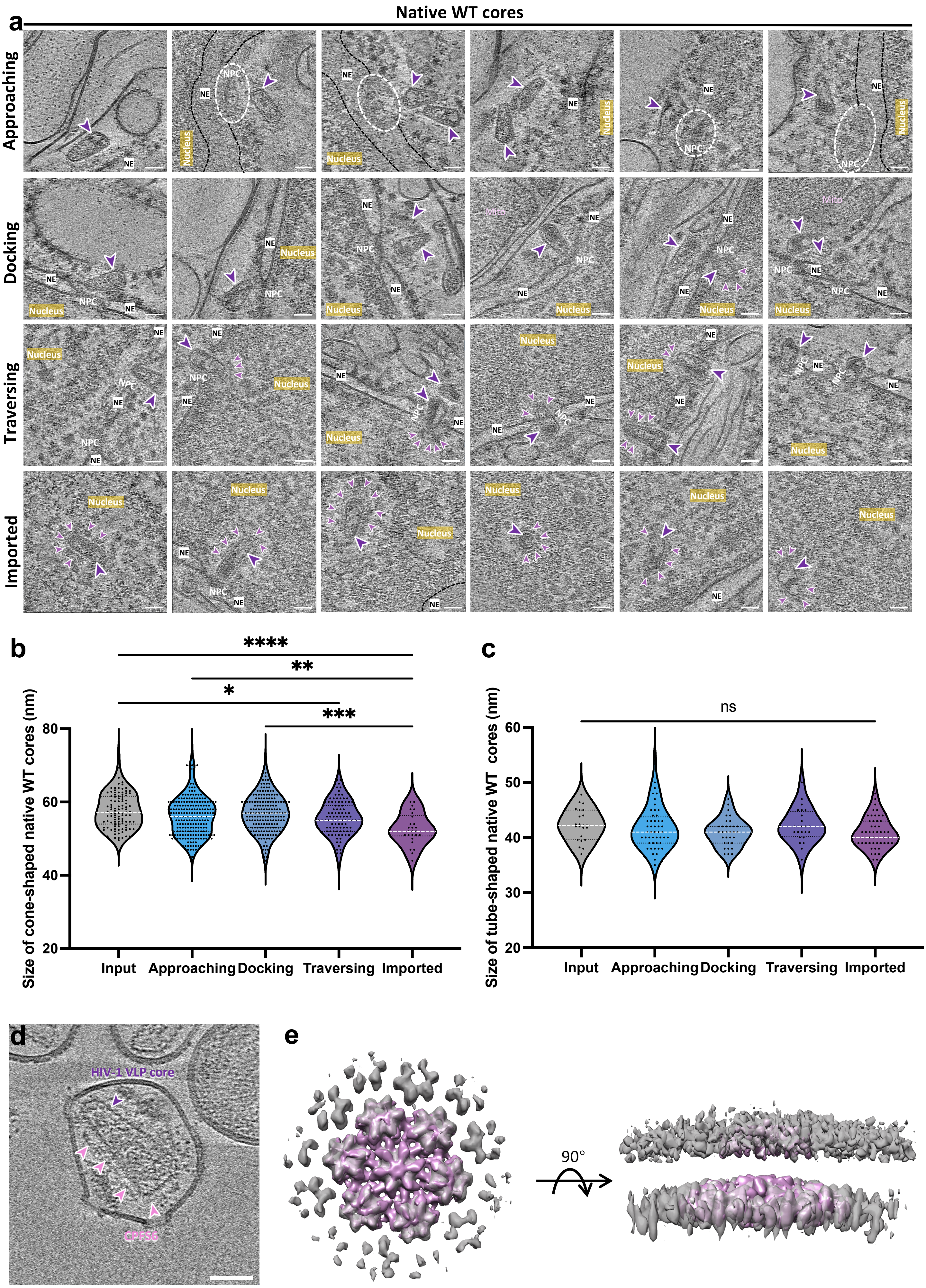

a
Native WT cores
Approaching
Docking
Traversing
Imported
NE
NPC
Nucleus
Mito
b
****
**
*
***
Size of cone-shaped native WT cores (nm)
80
60
40
20
Input
Approaching
Docking
Traversing
Imported
c
ns
Size of tube-shaped native WT cores (nm)
60
50
40
30
20
Input
Approaching
Docking
Traversing
Imported
d
HIV-1 VLP core
CPSF6
e
90°

**Extended Data Fig. 4 | Characterisation of native HIV-1 WT core nuclear import**. **a**, Gallery of native WT cores in multiple states during the nuclear import. Six representative tomographic slices from each state are showcased. Native WT cores are indicated by purple arrowheads, nuclear factors are indicated by light purple arrowheads, the nucleus, NE (indicated by black dashed lines in some cases), and NPC (indicated by white circles in the top view in some cases) are annotated accordingly. Scale bars = 50 nm. **b**, A violin plot of the statistical analysis on the size of cone-shaped native WT cores (width measured at the wide end) in each state. The size of imported cone-shaped native WT cores measures 52.65 ± 4.280 nm (SE = 0.8393, n = 26), the traversing measures 55.59 ± 4.594 nm (SE = 0.4617, n = 99), the docking measures 56.88 ± 4.996 nm (SE = 0.3673, n = 185), the approaching measures 56.12 ± 5.219 nm (SE = 0.3519, n = 220), and the input measures 57.71 ± 4.758 nm (SE = 0.4644, n = 105). White lines represent the medians, black lines represent the quartiles, and black dots represent individual cone-shaped native WT cores (One-way ANOVA test, **** = $p < 0.0001$, *** = $p < 0.001$, ** = $p < 0.01$, * = $p < 0.05$, only significant differences are shown, and annotated with asterisk). **c**, A violin plot of the statistical analysis on the size of tube-shaped native WT cores (width measured) in each state. The size of imported tube-shaped native WT cores measures 40.86 ± 2.750 nm (SE = 0.3850, n = 51), the traversing measures 42.17 ± 3.088 nm (SE = 0.6304, n = 24), the docking measures 41.00 ± 2.540 nm (SE = 0.4490, n = 32), the approaching measures 41.79 ± 3.848 nm (SE = 0.5554, n = 48), and the input measures 42.27 ± 2.875 nm (SE = 0.5996, n = 23). White lines represent the medians, black lines represent the quartiles, and black dots represent individual tube-shaped native WT cores (One-way ANOVA test, ns = no significance). **d**, A representative tomographic slice of a VLP treated with Streptolysin O (SLO) and incubated with purified CPSF6. HIV-1 cores and CPSF6 densities are labelled accordingly. **e**, The structural comparison between the CA hexamer structure (light purple) from imported native WT cores and the CA hexamer structure (grey, set to semi-transparent) from VLP cores incubated with purified CPSF6. The maps on the left are fitted and contoured at 3σ, while the top density on the right is highlighted with maps contoured at 0.5σ.

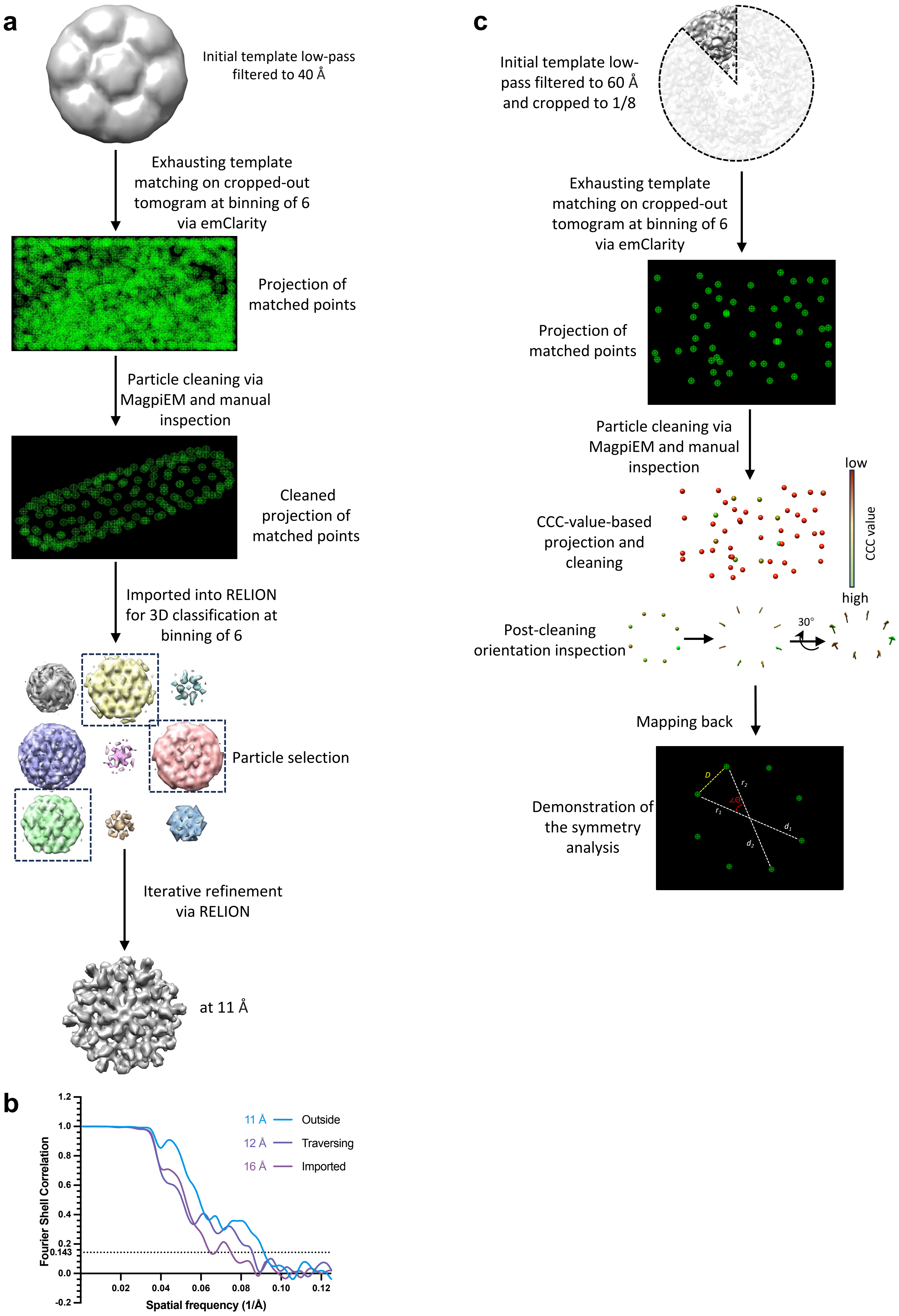

a
Initial template low-pass filtered to 40 Å
Exhausting template matching on cropped-out tomogram at binning of 6 via emClarity
Projection of matched points
Particle cleaning via MagpiEM and manual inspection
Cleaned projection of matched points
Imported into RELION for 3D classification at binning of 6
Particle selection
Iterative refinement via RELION
at 11 Å
b
Fourier Shell Correlation
Spatial frequency (1/Å)
11 Å Outside
12 Å Traversing
16 Å Imported
0.143
c
Initial template low-pass filtered to 60 Å and cropped to 1/8
Exhausting template matching on cropped-out tomogram at binning of 6 via emClarity
Projection of matched points
Particle cleaning via MagpiEM and manual inspection
CCC-value-based projection and cleaning
low
high
CCC value
Post-cleaning orientation inspection
30°
Mapping back
Demonstration of the symmetry analysis

**Extended Data Fig. 5 | Workflow of subtomogram averaging of native WT CA hexamers and symmetry analysis of NPCs**. **a**, The workflow used in this study for subtomogram averaging of CA hexamers. A low-pass filtered (40 Å) template was applied for the initial template matching using emClarity/1.5.0.2. The peak number was intentionally set to an excessive value in cropped tomograms to ensure an exhausting search. The matched particles were initially cleaned using MagpiEM, followed by manual inspection in Chimera to further remove false positives. The cleaned particles were then transferred into RELION/4.0 for 3D classification, good classes are selected for the following iterative refinement. **b**, Gold-standard Fourier shell correlation (FSC) curves of subtomogram averaged maps from imported (light purple), traversing (purple), and outside (approaching and docking combined, light blue) CA hexamers. The resolution is indicated at 0.143 FSC cut-off. **c**, The workflow for symmetry analysis of NPCs in this study. The initial NPC structure (EMD-11967) was low-pass filtered to 60 Å and cropped to $1/8^{th}$ of the original volume based on the 8-fold symmetry. The cropped map was then applied for the template matching via emClarity. The matched particles were initially cleaned by MagpiEM and then inspected in Chimera based on the cross-correlation value and orientation for further cleaning. The cleaned particles were then mapped back into the tomogram for the calculation of included angles between adjacent NPC subunits.

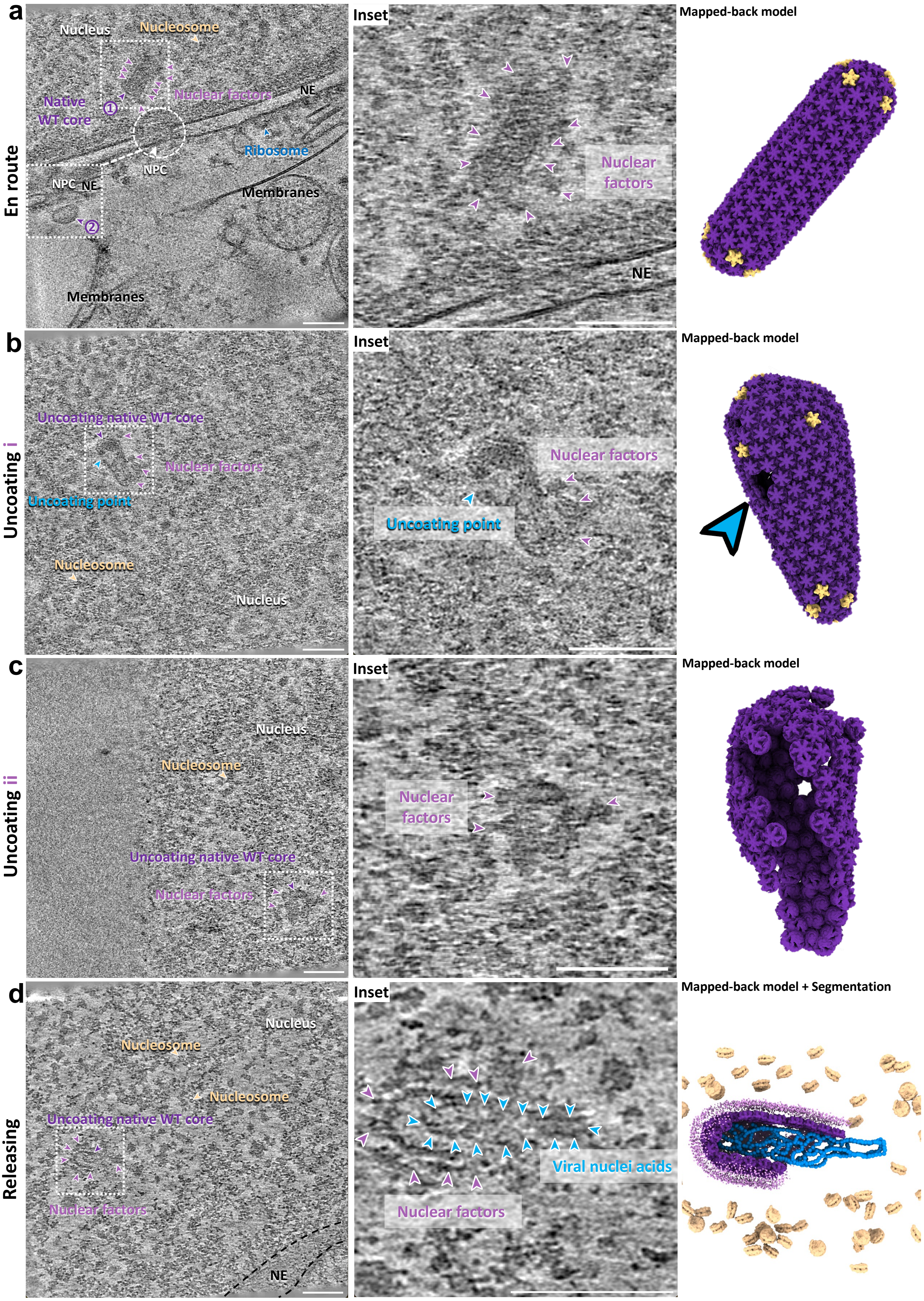

a
En route
Nucleus
Nucleosome
Native WT core
Nuclear factors
NE
NPC
Ribosome
Membranes
NPC NE
Membranes
Inset
Nuclear factors
NE
Mapped-back model
b
Uncoating i
Uncoating native WT core
Nuclear factors
Uncoating point
Nucleosome
Nucleus
Inset
Nuclear factors
Uncoating point
Mapped-back model
c
Uncoating ii
Nucleus
Nucleosome
Uncoating native WT core
Nuclear factors
Inset
Nuclear factors
Mapped-back model
d
Releasing
Nucleus
Nucleosome
Nucleosome
Uncoating native WT core
Nuclear factors
NE
Inset
Viral nuclei acids
Nuclear factors
Mapped-back model + Segmentation

**Extended Data Fig. 6 | Visualisation of native HIV-1 WT core nuclear trafficking. a,** A representative tomographic slice of a correlatively-acquired tomogram containing native WT cores. Two native WT cores were identified and indicated by purple arrowheads and numbered: No.1, an imported tube-shaped native WT core being transported in the nucleus with discernable surrounding densities; No.2, a docked cone-shaped native WT core on the NPC, shown on another slice of the same tomogram (white framed). The zoomed-in view of the native HIV-1 WT core en route is depicted in the middle panel. The mapped-back model of this native WT core is depicted in the right panel, CA pentamers are highlighted in gold colour. **b-d,** Representative tomographic slices of correlatively-acquired tomograms of native WT cores uncoating in the nucleus: incipient (**b**), half-way (**c**), and nucleic acid-releasing (**d**). In **b**, the uncoating point is indicated by the light blue arrowhead. In **c,** half of the capsid could not be detected by template matching and is apparently missing. In **d**, nucleic acids can be seen releasing from the remaining capsid shell; nucleic acid density is traced as indicated by light blue arrowheads. Mapped-back models and segmented volumes are illustrated in the right panel. The NPC, ribosomes, and nucleosomes, surrounding nuclear factors, and prominent linker DNA between nucleosomes are labelled. The nucleus, nuclear envelope (NE) and membranes are annotated accordingly. Scale bar = 100 nm.

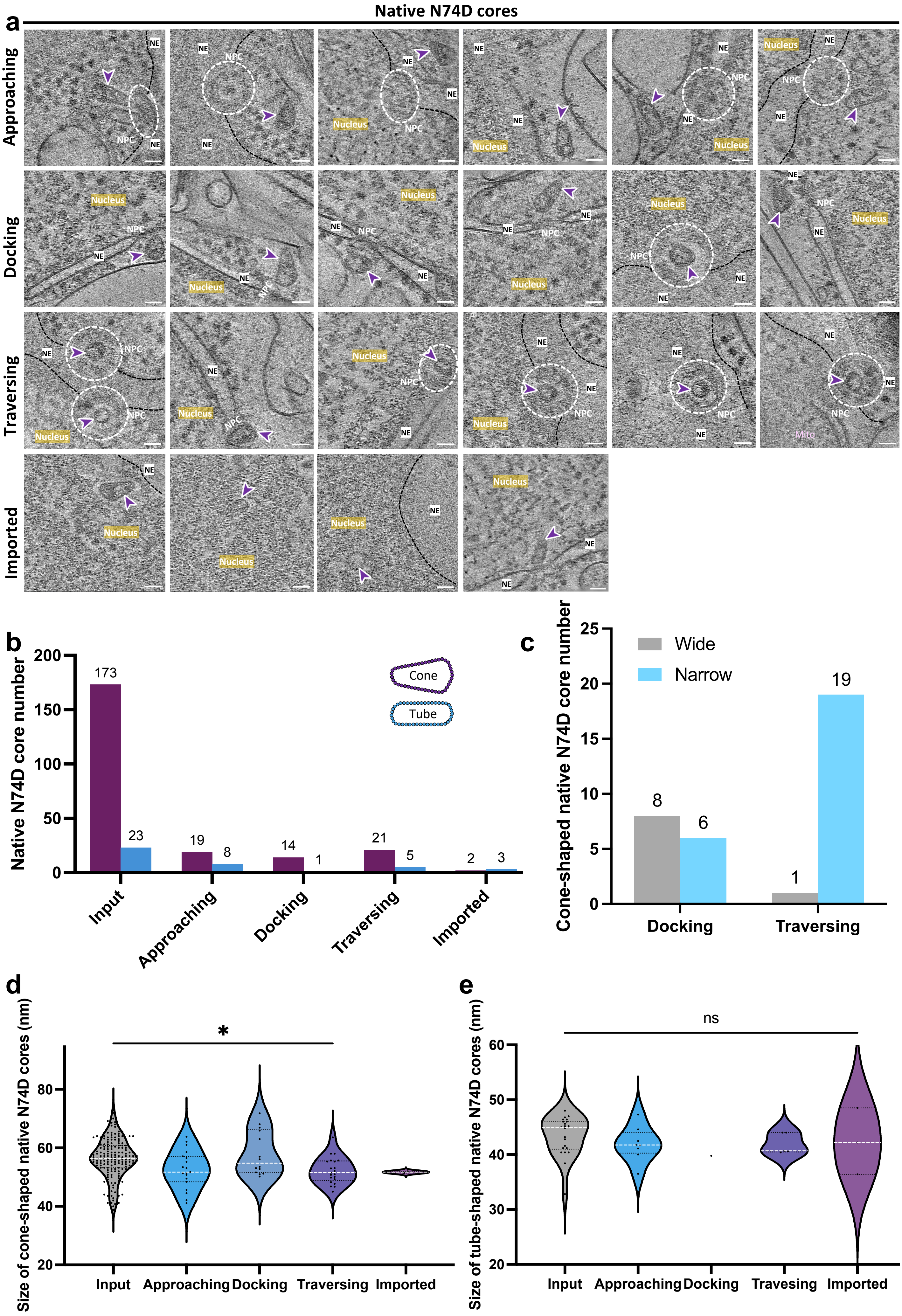

a
Native N74D cores
Approaching
Docking
Traversing
Imported
NE
NPC
Nucleus
Mito
b
Native N74D core number
Cone
Tube
173
23
19
8
14
1
21
5
2
3
Input
Approaching
Docking
Traversing
Imported
c
Cone-shaped native N74D core number
Wide
Narrow
8
6
1
19
Docking
Traversing
d
Size of cone-shaped native N74D cores (nm)
*
Input
Approaching
Docking
Traversing
Imported
e
Size of tube-shaped native N74D cores (nm)
ns
Input
Approaching
Docking
Travesing
Imported

**Extended Data Fig. 7 | Characterisation of native HIV-1 N74D core nuclear import**. **a**, Gallery of native N74D cores in multiple states during nuclear import. Six representative tomographic slices are shown for each state, except for the imported state, where four slices are displayed. Native N74D cores are indicated by purple arrowheads, the nucleus, NE (indicated by black dashed lines in some cases), and NPC (indicated by white circles in the top view in some cases) are annotated accordingly. Scale bars = 50 nm. **b**, A bar chart illustrating the composition of native N74D core shapes in each state. Cone-shaped cores are in purple and tube-shaped cores are in blue (Chi-square test for all, $p = 0.0739$). **c**, A bar chart showing the orientation distribution of cone-shaped native N74D cores in docking and traversing states, with the wide end in first (grey) and narrow end in first (light blue) (Fisher's exact test, $p = 0.0012$). **d**, A violin plot of the statistical analysis on the size of cone-shaped native N74D cores (width measured at the wide end) in each state. The size of imported cone-shaped native N74D cores measures $51.70 \pm 0.4243$ nm (SE = 0.3000, n = 2), the traversing measures $52.23 \pm 4.536$ nm (SE = 0.9899, n = 21), the docking measures $58.01 \pm 7.504$ nm (SE = 2.006, n = 14), the approaching measures $52.48 \pm 6.484$ nm (SE = 1.487, n = 19), and the input measures $56.46 \pm 6.644$ nm (SE = 0.5052, n = 173). White lines represent the medians, black lines represent the quartiles, and black dots represent individual cone-shaped native N74D cores (One-way ANOVA test, * = $p < 0.05$, only significant differences are shown, and annotated with asterisk). **e**, A violin plot of the statistical analysis on the size of tube-shaped native N74D cores in each state. The size of imported tube-shaped native N74D cores measures $42.37 \pm 6.052$ nm (SE = 3.494, n = 3), the traversing measures $41.94 \pm 1.884$ nm (SE = 0.8424, n = 5), the docking measures 39.80 (n =1), the approaching measures $41.94 \pm 3.168$ nm (SE = 1.120, n = 8), and the input measures $43.40 \pm 3.590$ nm (SE = 0.7655, n = 22). White lines represent the medians, black lines represent the quartiles, and black dots represent individual tube-shaped native N74D cores (One-way ANOVA test, ns = no significance).

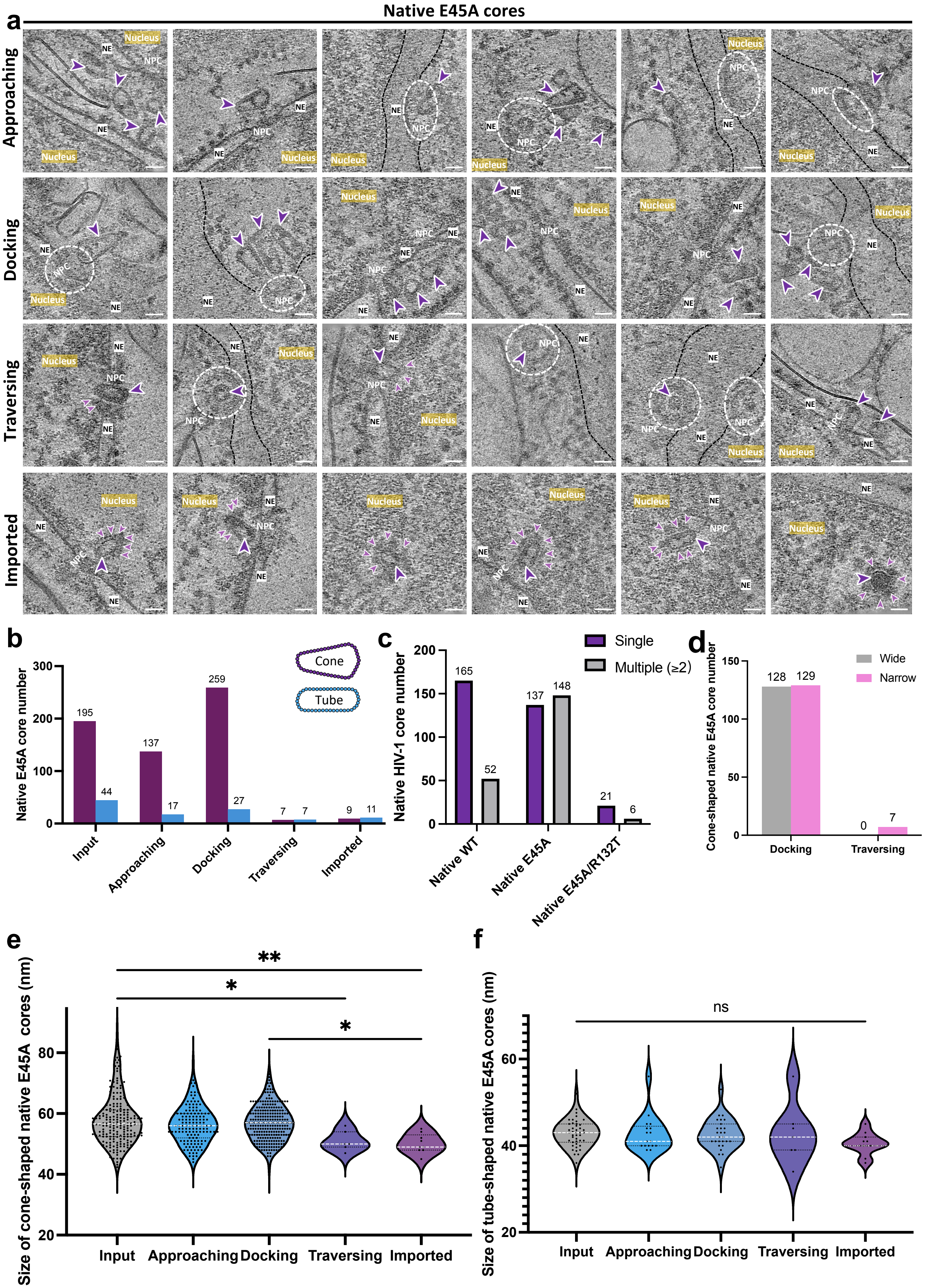

Native E45A cores
a
Approaching
Docking
Traversing
Imported
Nucleus
NE
NPC
b
Native E45A core number
Cone
Tube
195
44
137
17
259
27
7
7
9
11
Input
Approaching
Docking
Traversing
Imported
c
Native HIV-1 core number
Single
Multiple (≥2)
165
52
137
148
21
6
Native WT
Native E45A
Native E45A/R132T
d
Cone-shaped native E45A core number
Wide
Narrow
128
129
0
7
Docking
Traversing
e
Size of cone-shaped native E45A cores (nm)
**
*
*
Input
Approaching
Docking
Traversing
Imported
f
Size of tube-shaped native E45A cores (nm)
ns
Input
Approaching
Docking
Traversing
Imported

**Extended Data Fig. 8 | Characterisation of native E45A core**. **a**, Gallery of native E45A cores in multiple states during the nuclear import. Six representative tomographic slices from each state are showcased. Native E45A cores are indicated by purple arrowheads, nuclear factors are indicated by light purple arrowheads, the nucleus, NE (indicated by black dashed lines in some cases), and NPC (indicated by white circles in the top view in some cases) are annotated accordingly. Scale bars = 50 nm. **b**, A bar chart illustrating the composition of native E45A core shapes in each state. Cone-shape cores are in purple and tube-shaped cores are in blue (Chi-square test, $p < 0.0001$). **c**, A bar chart showing the distribution of native WT, E45A, and E45A/R132T cores docking at a single NPC. Single: only one core; Multiple: ≥ two cores (Chi-square test for all, and Fisher's exact test for E45A and WT, $p < 0.0001$). **d**, A bar chart showing the orientation distribution of cone-shaped native E45A cores in docking and traversing states, with the wide end in first (grey) and narrow end in first (pink) (Fisher's exact test, $p = 0.0147$). **e**, A violin plot of the statistical analysis on the size of cone-shaped native E45A cores (width measured at the wide end) in each state. The size of imported cone-shaped native E45A cores measures $50.11 \pm 3.180$ nm (SE = 1.060, n = 9), the traversing measures $50.86 \pm 3.078$ nm (SE = 1.164, n = 7), the docking measures $56.86 \pm 5.791$ nm (SE = 0.3598, n = 259), the approaching measures $56.32 \pm 6.054$ nm (SE = 0.5172, n = 137), and the input measures $57.89 \pm 8.162$ nm (SE = 0.5845, n = 195). White lines represent the medians, black lines represent the quartiles, and black dots represent individual cone-shaped native E45A cores (One-way ANOVA test, ** = $p < 0.01$, * = $p < 0.05$, only significant differences are shown). **f**, A violin plot of the statistical analysis on the size of tube-shaped native E45A cores (width measured) in each state. The size of imported tube-shaped native E45A cores measures $40.45 \pm 2.505$ nm (SE = 0.7551, n = 11), the traversing measures $42.71 \pm 6.921$ nm (SE = 2.616, n = 7), the docking measures $42.63 \pm 3.596$ nm (SE = 0.6921, n = 27), the approaching measures $42.76 \pm 4.176$ nm (SE = 1.013, n = 17), and the input measures $42.84 \pm 3.116$ nm (SE = 0.4751, n = 43). White lines represent the medians, black lines represent the quartiles, and black dots represent individual tube-shaped native E45A cores (One-way ANOVA test, ns = no significance).

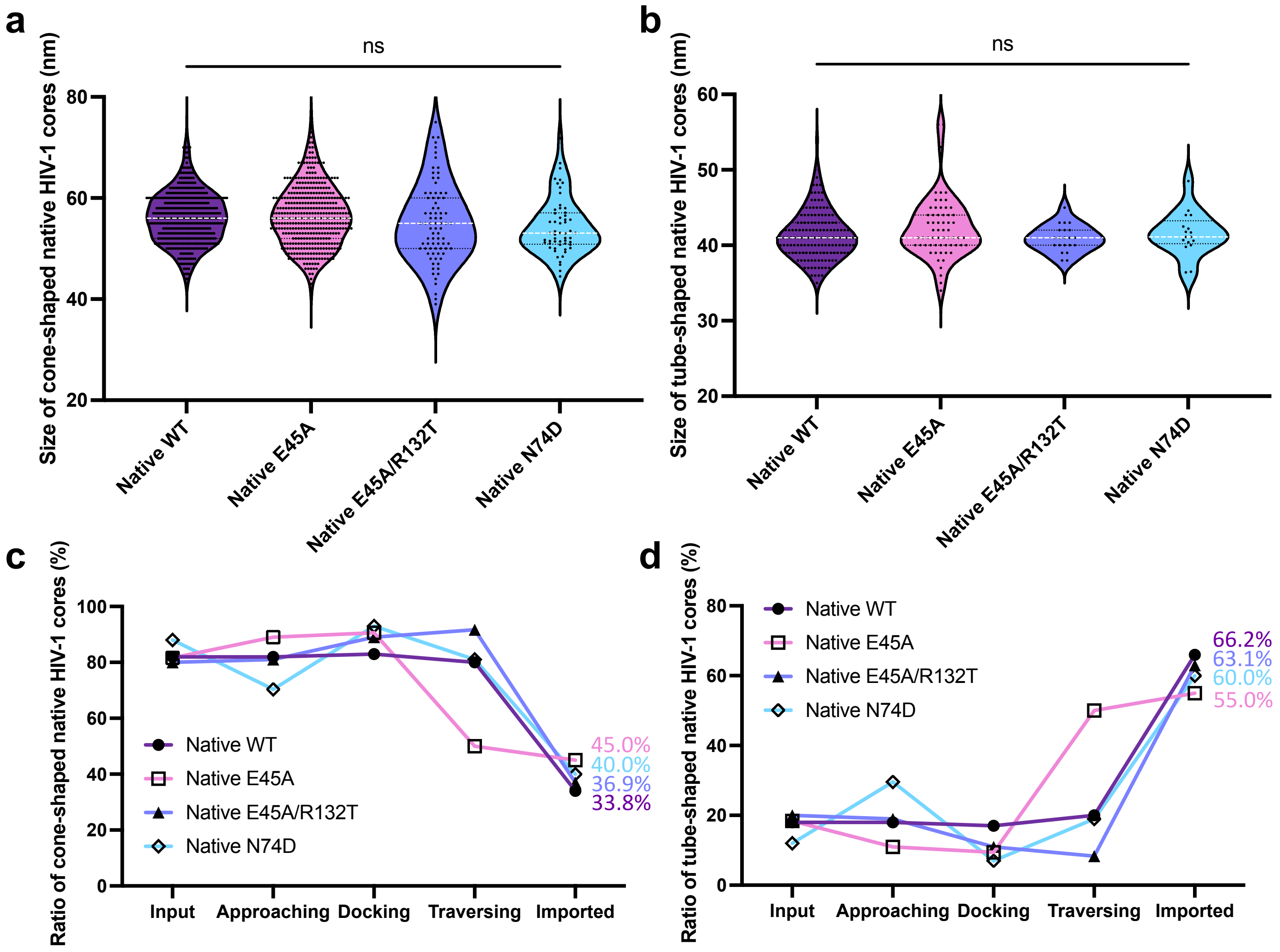


**Extended Data Fig. 9 | Characterisation of the nuclear import of native HIV-1 cores.** **a**, A violin plot of the statistical analysis on the size of cone-shaped HIV-1 cores (width measured at the wide end) from all native samples incubated with PS-CEM cells. The size of cone-shaped native WT cores measures 56.11 ± 5.053 nm (SE = 0.2193, n = 530), the E45A measures 56.43 ± 2.919 nm (SE = 0.2916, n = 412), the E45A/R132T measures 55.10 ± 7.802 nm (SE = 0.8668, n = 79), and the N74D measures 54.36 ± 5.587 nm (SE = 0.7466, n = 56). White lines represent the medians, black lines represent the quartiles, and black dots represent individual cone-shaped native cores (One-way ANOVA test, ns = no significance). **b**, A violin plot of the statistical analysis on the size of tube-shaped cores (width measured) from all native samples incubated with PS-CEM cells. The size of tube-shaped native WT cores measures 41.38 ± 3.157 nm (SE = 0.2535, n = 155), the E45A measures 42.29 ± 4.071 nm (SE = 0.5170, n = 62), the E45A/R132T measures 40.90 ± 1.758 nm (SE = 0.3836, n = 23), and the N74D measures 41.52 ± 2.871 nm (SE = 0.6962, n = 17). White lines represent the medians, black lines represent the quartiles, and black dots represent individual tube-shaped native HIV-1 cores (One-way ANOVA test, ns = no significance). **c**, A line chart illustrating the percentage of cone-shaped HIV-1 cores in each state of all native samples incubated with PS-CEM cells. **d**, A line chart illustrating the percentage of tube-shaped HIV-1 cores in each state of all native samples incubated with PS-CEM cells.

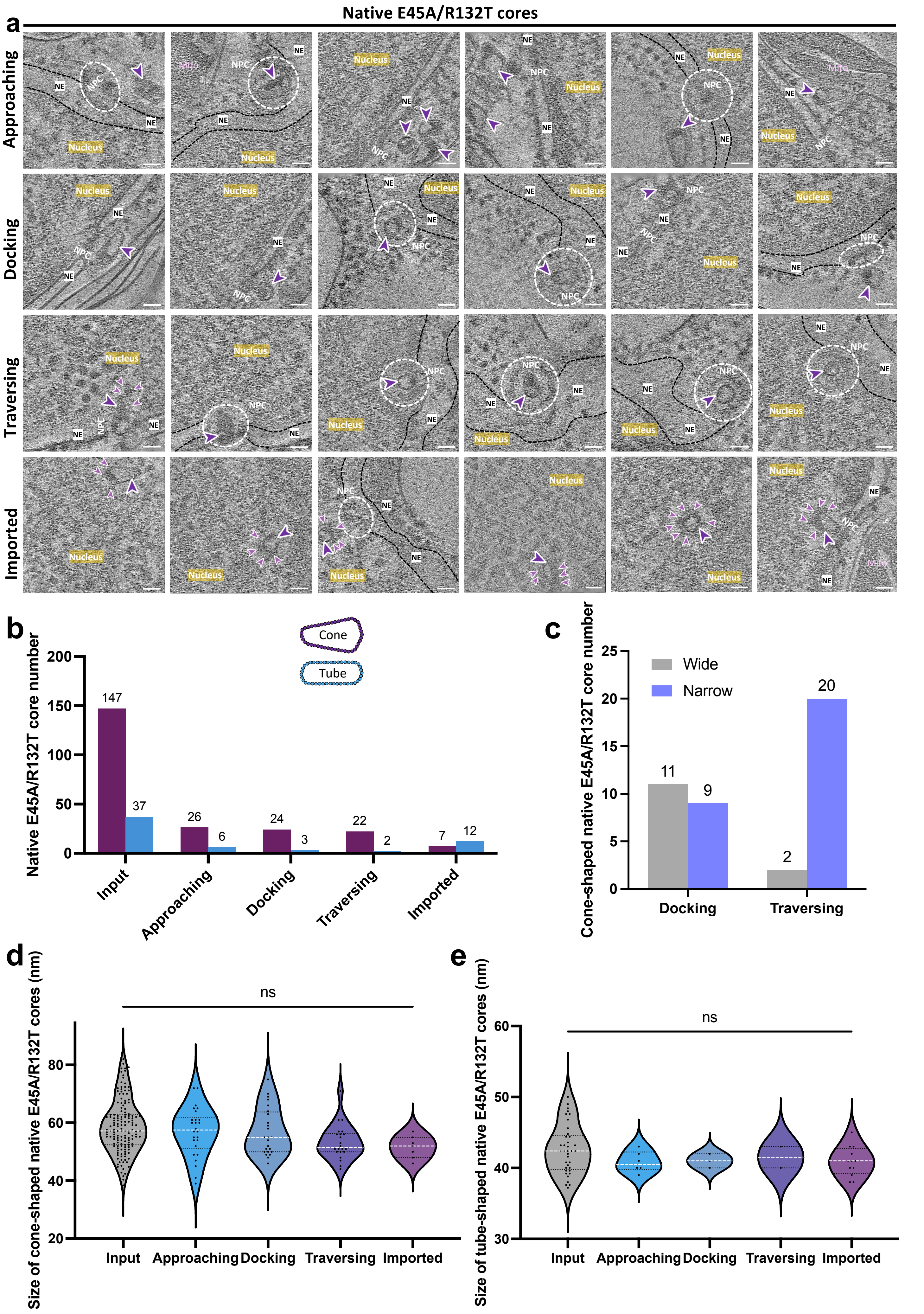

a
Native E45A/R132T cores
Approaching
Docking
Traversing
Imported
Nucleus
NE
NPC
Mito
b
Cone
Tube
Native E45A/R132T core number
200
150
100
50
0
147
37
26
6
24
3
22
2
7
12
Input
Approaching
Docking
Traversing
Imported
c
Cone-shaped native E45A/R132T core number
25
20
15
10
5
0
Wide
Narrow
11
9
2
20
Docking
Traversing
d
Size of cone-shaped native E45A/R132T cores (nm)
ns
80
60
40
20
Input
Approaching
Docking
Traversing
Imported
e
Size of tube-shaped native E45A/R132T cores (nm)
ns
60
50
40
30
Input
Approaching
Docking
Traversing
Imported

**Extended Data Fig. 10 | Characterisation of native E45A/R132T core nuclear import**. **a**, Gallery of native E45A/R132T cores in multiple states during nuclear import. Six representative tomographic slices from each state are shown. Native E45A/R132T cores are indicated by purple arrowheads, nuclear factors are indicated by light purple arrowheads, the nucleus, NE (indicated by black dashed lines in some cases), and NPC (indicated by white circles in the top view in some cases) are annotated accordingly. Scale bars = 50 nm. **b**, A bar chart illustrating the composition of native E45A/R132T core shapes in each state. Cone-shape cores are in purple and tube-shaped cores in blue (Chi-square test for all, $p < 0.0001$). **c**, A bar chart showing the orientation distribution of cone-shaped native E45A/R132T cores in docking and traversing states, with the wide end in first (grey) and narrow end in first (light purple) (Fisher's exact test, $p = 0.0022$). **d**, A violin plot of the statistical analysis on the size of cone-shaped native E45A/R132T cores (width measured at the wide end) in each state. The size of imported cone-shaped native E45A/R132T cores measures 51.57 $\pm$ 3.867 nm (SE = 1.462, n = 7), the traversing measures 53.32 $\pm$ 6.027 nm (SE = 1.285, n = 22), the docking measures 56.96 $\pm$ 8.164 nm (SE = 1.666, n = 24), the approaching measures 56.88 $\pm$ 8.454 nm (SE = 1.658, n = 26), and the input measures 58.66 $\pm$ 9.166 nm (SE = 0.7560, n = 147). White lines represent the medians, black lines represent the quartiles, and black dots represent individual cone-shaped native E45A/R132T cores (One-way ANOVA test, ns = no significance). **e**, A violin plot of the statistical analysis on the size of tube-shaped native E45A/R132T cores (width measured) in each state. The size of imported tube-shaped native E45A/R132T cores measures 40.92 $\pm$ 2.109 nm (SE = 0.6088, n = 12), the traversing measures 41.50 $\pm$ 2.121 nm (SE = 1.500, n = 2), the docking measures 41.00 $\pm$ 1.000 nm (SE = 0.5774, n = 3), the approaching measures 40.83 $\pm$ 1.472 nm (SE = 0.6009, n = 6), and the input measures 42.63 $\pm$ 3.460 nm (SE = 0.5688, n = 37). White lines represent the medians, black lines represent the quartiles, and black dots represent individual tube-shaped native E45A/R132T cores (One-way ANOVA test, ns = no significance).

**Table 1 |** Shape distribution of VLP HIV-1 cores in P-CEM cells in multiple states.

| | Unassociated | Approaching | Docking | Passage | Imported |
|---|---|---|---|---|---|
| Cone-shaped | 17 | 31 | 50 | 33 | 2 |
| Tube-shaped | 40 | 62 | 72 | 117 | 43 |
| Total | 57 | 93 | 122 | 150 | 45 |

**Table 2 |** Shape distribution of native HIV-1 WT cores in P-CEM cells in multiple states

| | Approaching | Docking | Passage | Imported |
|---|---|---|---|---|
| Cone-shaped | 61 | 48 | 14 | 2 |
| Tube-shaped | 9 | 9 | 4 | 8 |
| Total | 70 | 57 | 18 | 10 |

**Table 3 |** Shape distribution of native HIV-1 WT cores in PS-CEM cells in multiple states

| | Approaching | Docking | Passage | Imported |
|---|---|---|---|---|
| Cone-shaped | 220 | 185 | 99 | 26 |
| Tube-shaped | 48 | 32 | 24 | 51 |
| Total | 268 | 217 | 123 | 77 |

**Table 4 |** Shape distribution of native HIV-1 E45A cores in PS-CEM cells in multiple states

| | Approaching | Docking | Passage | Imported |
|---|---|---|---|---|
| Cone-shaped | 137 | 259 | 7 | 9 |
| Tube-shaped | 17 | 27 | 7 | 11 |
| Total | 154 | 286 | 14 | 20 |

**Table 5 |** Shape distribution of native HIV-1 E45A/R132T cores in PS-CEM cells in multiple states

| | Approaching | Docking | Passage | Imported |
|---|---|---|---|---|
| Cone-shaped | 26 | 24 | 22 | 7 |
| Tube-shaped | 6 | 3 | 2 | 12 |
| Total | 32 | 27 | 24 | 19 |

**Table 6 |** Shape distribution of native HIV-1 N74D cores in PS-CEM cells in multiple states

| | Approaching | Docking | Passage | Imported |
|---|---|---|---|---|
| Cone-shaped | 19 | 14 | 21 | 2 |
| Tube-shaped | 8 | 1 | 5 | 3 |
| Total | 27 | 15 | 26 | 5 |

**Table 7 |** Orientation distribution of cone-shaped HIV-1 VLP cores in docking and passage states in P-CEM cells

| | Docking | Passage |
|---|---|---|
| Wide end | 35 | 9 |
| Narrow end | 15 | 24 |

**Table 8 |** Orientation distribution of cone-shaped native HIV-1 WT cores in docking and passage states in PS-CEM cells

| | Docking | Passage |
|---|---|---|
| Wide end | 93 | 3 |
| Narrow end | 87 | 89 |

**Table 9 |** Orientation distribution of cone-shaped native HIV-1 E45A cores in docking and passage states in PS-CEM cells

| | Docking | Passage |
|---|---|---|
| Wide end | 128 | 0 |
| Narrow end | 129 | 7 |

**Table 10 |** Orientation distribution of cone-shaped native E45A/R132T cores in docking and passage states in PS-CEM cells

| | Docking | Passage |
|---|---|---|
| Wide end | 11 | 2 |
| Narrow end | 9 | 20 |

**Table 11 |** Orientation distribution of cone-shaped native HIV-1 E45A/R132T cores in docking and passage states in PS-CEM cells

| | Docking | Passage |
|---|---|---|
| Wide end | 8 | 1 |
| Narrow end | 6 | 19 |

**Table 12 |** Cryo-FIB lamella preparation

| **Method** | Correlative milling | Correlative milling | Correlative lift-out |
|---|---|---|---|
| Microscope | Plasma FIB Arctis | Conventional FIB Aquilos 2 | Conventional FIB Aquilos 2 |
| Voltage (keV) | 30 | 30 | 30 |
| Ion beam source | Argon | Gallium | Gallium |
| Sputtering coating prior to milling (seconds) | 12 | No | 12 |
| GIS coating time (second) | 50 | 30 | 30 |
| Bulk milling current | N/A | N/A | 5-7 nA |
| Milling current | 0.74-2 nA | 0.1-0.5 nA | 0.1-0.5 nA |
| Polishing current | 60 pA | 30 pA | 30 pA |
| Sputtering coating post polishing (seconds) | No | No | No |
| Fluorescence microscope | iFLM (100 ×) | METEOR (50 ×) | iFLM (20 ×) |
| Number of lamellae | 81 | 398 | 5 |

**Table 13 |** Cryo-ET data collection

| **Sample** | Lamellae of VLP cores | Lamellae of native WT cores | Lamellae of native E45A cores | Lamellae of native E45A/R132T cores | Lamellae of native N74D cores | All isolated cores on grids | HIV-1 virions | Lamellae of CEM cells | CPSF6 bound to perforated VLP |
|---|---|---|---|---|---|---|---|---|---|
| Microscope | FEI Titan Krios G3/G4 | FEI Titan Krios G3 | FEI Titan Krios G3 | FEI Titan Krios G3 | FEI Titan Krios G3 | FEI Titan Krios G2 | FEI Titan Krios G2 | FEI Titan Krios G2 | FEI Titan Krios G2 |
| Voltage (keV) | 300 | 300 | 300 | 300 | 300 | 300 | 300 | 300 | 300 |
| Detector | Falcon 4i | Falcon 4i | Falcon 4i | Falcon 4i | Falcon 4i | Gatan K3 | Gatan K3 | Gatan K3 | Falcon 4i |
| Energy-filter | Selectris X | Selectris X | Selectris X | Selectris X | Selectris X | Gatan BioQuantum | Gatan BioQuantum | Gatan BioQuantum | Selectris X |
| Slit width (eV) | 10 | 10 | 10 | 10 | 10 | 20 | 20 | 20 | 20 |
| Super-resolution mode | No | No | No | No | No | Yes | No | Yes | No |
| Physical pixel size (Å/pixel) | 1.903/1.94/2 | 1.903/1.94 | 1.903/1.94 | 1.903 | 1.94 | 0.831 | 1.34 | 2.18 | 1.34 |
| Defocus range (µm) | -3 to -5, increment 0.3 | -3 to -5, increment 0.3 | -3 to -5, increment 0.3 | -3 to -5, increment 0.3 | -3 to -5, increment 0.3 | -3 to -4, increment 0.3 | -1.5 to -3, increment 0.3 | -3 to -5, increment 0.3 | -1.5 to -3, increment 0.3 |
| Acquisition scheme | Dose-Symmetric, -52º/52º, -54º/54º -60º/60º, 2º step, group 2 | Dose-Symmetric, -52º/52º, -54º/54º, 2º step, group 2 | Dose-Symmetric, -54º/54º, 2º step, group 2 | Dose-Symmetric, -54º/54º, 2º step, group 2 | Dose-Symmetric, -54º/54º, 2º step, group 2 | Single-shot micrograph | Dose-Symmetric, -60º/60º, 3º step, group 3 | Dose-Symmetric, -54º/54º, 3º step, group 3 | Dose-Symmetric, -60º/60º, 3º step, group 3 |
| Total dose (electrons/Å$^2$) | 159/137.5/152.5 | 137.5/132.5 | 137.5/132.5 | 137.5 | 137.5 | 22 | 123 | 37 | 120 |
| Number of frames | 10/EER | 10 | 10 | 10 | 10 | 59 | 10 | 10 | 10 |
| Number of lamellae | 125 | 185 | 72 | 32 | 29 | N/A | N/A | 10 | N/A |
| Number of tomograms/micrographs | 269 | 759 | 322 | 122 | 179 | 10,000 | 58 | 47 | 100 |

**Table 14 |** Structural determination of native HIV-1 WT CA hexamers

| **States of native HIV-1 WT cores** | Outside | Passage | Imported |
|---|---|---|---|
| Particle number | 11,915 | 5,545 | 7,825 |
| Final resolution by gold-standard FSC cut (Å) | 11.0 | 11.7 | 15.8 |
